# The multifunction *Coxiella* effector Vice stimulates macropinocytosis and interferes with the ESCRT machinery

**DOI:** 10.1101/2024.03.13.584753

**Authors:** Arthur Bienvenu, Melanie Burette, Franck Cantet, Manon Gourdelier, Jitendriya Swain, Chantal Cazevieille, Tatiana Clemente, Arif Sadi, Claire Dupont, Manon Le Fe, Nicolas Bonetto, Benoit Bordignon, Delphine Muriaux, Stacey Gilk, Matteo Bonazzi, Eric Martinez

## Abstract

Intracellular bacterial pathogens divert multiple cellular pathways to establish their niche and persist inside their host. *Coxiella burnetii*, the causative agent of Q fever, secretes bacterial effector proteins via its Type 4 secretion system to generate a *Coxiella*-containing vacuole (CCV). Manipulation of lipid and protein trafficking by these effectors is essential for bacterial replication and virulence. Here, we have characterized the lipid composition of CCVs and discovered that the effector Vice interacts with phosphoinositides and membranes enriched in phosphatidylserine (PS) and lysobisphosphatidic acid (LBPA). Remarkably, eukaryotic cells ectopically expressing Vice present compartments that resemble early CCVs in both morphology and composition. We discovered that the biogenesis of these compartments relies on the double function of Vice. The effector protein initially localizes at the plasma membrane of eukaryotic cells where it triggers the internalization of large vacuoles by macropinocytosis. Then, Vice stabilizes these compartments by perturbing the ESCRT machinery and inhibiting the formation of intraluminal vesicles (ILVs). Collectively, our results reveal that Vice is an essential *C. burnetii* effector protein capable of hijacking two major cellular pathways to shape the bacterial replicative niche.

**Significance statement:** *Coxiella burnetii* is a unique bacterial pathogen that secretes more than a hundred effector proteins to manipulate cellular processes and establish a replicative niche, the *Coxiella*-containing vacuole (CCV). Our study identified host cell lipids that are actively recruited by the bacterium to the CCV. Using a library of effector mutants, we identified the protein Vice (for Vacuole-inducing *Coxiella* effector) as the first bacterial effector capable of interacting with lysobisphosphatydic acid-enriched membranes and accumulating this lipid to CCVs. We show that Vice is also capable of stimulating macropinocytosis and inhibiting the ESCRT machinery. Together, our data show how a single bacterial effector can manipulate different cellular processes to favor the biogenesis of a bacterial pathogen’s niche.

## Introduction

Intracellular bacterial pathogens have evolved a variety of strategies to survive and replicate inside infected cells. While intra-cytosolic bacteria actively escape phagosomes and establish their replicative niche in the cytoplasm, intravacuolar bacteria remain confined within host-derived compartments and escape degradation by subverting membrane trafficking to generate pathogen-specific replicative niches(1).

The obligate intracellular bacterium *Coxiella burnetii* is a highly infectious pathogen responsible for the re-emerging zoonosis Q fever, a debilitating flu-like disease with severe health and economic burdens(2). Upon internalization by host cells, *C. burnetii* remains confined within membrane-bound compartments that follow the degradative endocytic pathway. Once in an acidic and degradative environment, the bacterium activates its Type 4b Secretion System (T4SS) to translocate over 140 effector proteins into the host cell cytoplasm(3). These effector proteins subvert multiple host cell functions to ensure the biogenesis of *Coxiella*-containing vacuoles (CCVs). A remarkable feature of CCV biogenesis is the rapid expansion of these compartments that precedes bacterial replication. Thus, at the early stages of infection, CCVs appear as large, spacious intracellular compartments harboring few bacteria. Later during infection, bacterial replication ensues and CCVs are rapidly filled with bacteria. While the role of host proteins subverted by *C. burnetii* during infections is well investigated, lipids are emerging as important players in *C. burnetii* intracellular replication and in CCV biogenesis. For example, CvpB (or Cig2) is a *C. burnetii* lipid-interacting effector (LIE) that triggers an increase of phosphatidylinositol 3-phosphate (PI(3)P) at CCVs and favors the homotypic fusion of CCVs(4). Notably, the characterization of CvpB revealed an unexpected lipid composition of CCVs and demonstrated how defective CCV biogenesis severely impacts *C. burnetii* virulence(5).

Lipids constitute an essential component of the “membrane code”(6), regulating signal transduction, cytoskeleton architecture and membrane trafficking. Specific lipid-modifying enzymes control lipid spatial and temporal distribution allowing precise and local modulation of essential processes including endocytosis and phagocytosis, membrane traffic and autophagy(7). Phosphatidylinositol can be phosphorylated on three carbons to give rise to 7 different phospholipids essential to orchestrate cellular trafficking processes(7).

Several bacterial effector proteins manipulate PI metabolism to favor bacterial intracellular replication: *Mycobacterium tuberculosis* secretes the PI(3)P phosphatase SapM(8) and the broad range phosphatase MptpB(9); *Legionella pneumophila* secretes the effector SidP(10) to hydrolyse PI(3)P and PI(3,5)P_2_, and effectors SidC, SidM, LidA, LpnE and RidL to use PI(3)P and/or PI(4)P as membrane anchors. Additionally, *Salmonella enterica* serovar Typhimurium increases PI(3)P in infected cells through the activity of the SPI-1 Type 3 secretion system (T3SS) substrate SopB(11, 12), a pleiotropic PI polyphosphatase. Besides promoting *Salmonella* uptake by macropinocytosis(13), SopB also maintains high levels of PI(3)P at the surface of *Salmonella*-containing vacuoles (SCVs) to form large SCVs.(12).

Given the importance of host lipids in *C. burnetii* intracellular development and virulence, we have further investigated the lipid composition of CCVs using lipid-binding sensors and antibodies and screened *C. burnetii* mutants to identify bacterial proteins that could be involved in the manipulation of lipid trafficking during infection. Our results show that PI(3,5)P_2_ and PS are passively recruited at CCV membranes, whereas PI(3)P, PI(4)P and lysobisphosphatydic acid (LBPA) are enriched at CCV membranes in a T4SS-dependent manner. We also show that LBPA localization at CCVs depends on the activity of Vice (for Vacuole-inducing *Coxiella* effector, encoded by the gene *cbu2007*), a *C. burnetii* effector protein responsible for the initial CCV expansion and stabilization during infection. Intriguingly, the ectopic expression of Vice in non-phagocytic cells triggers macropinocytosis, leading to the formation of Vice-induced compartments (VICs), large intracellular vacuoles, reminiscent in morphology and composition of early CCVs and bearing multivesicular bodies (MVB) markers. At later time points of expression, Vice perturbs the ESCRT machinery, thereby stabilizing VICs, possibly by perturbing the formation of intraluminal vesicles (ILVs). We propose that Vice is a multifunction *C. burnetii* effector protein essential for the onset of CCVs biogenesis.

## Results

### PI(3)P, PI(4)P and LBPA are enriched at CCV membranes in a T4SS-dependent manner

To determine the lipid composition of CCVs, a panel of well-established lipid-binding sensors (SI appendix, Table S1) was cloned into mCherry- or RFP-expressing vectors and used to monitor lipid localization by automated confocal live cell imaging of U2OS cells infected with GFP-tagged wt *C. burnetii* for 2, 4 and 6 days. U2OS cells transfected with a plasmid encoding mCherry alone were used as control. Due to the lack of specific sensors, LBPA localization was monitored in U2OS cells infected with *C. burnetii* as above using anti-LBPA antibodies following fixation. The presence of each lipid at CCVs was visually scored from an average of 100 cells per condition (Fig. 1A). mCherry alone was never observed accumulating at CCV membranes (Fig. 1B). At all time points of infection, over 90% of infected cells presented CCVs positive for PS, a typical marker of late endosomal/lysosomal compartments (Fig. 1A, B). Interestingly however, only half of the infected cells presented CCVs that were positive for PI(3,5)P_2_, another major lipid marker of late endosomes and lysosomes (Fig. 1A, B). This is in line with our previous observation that the *C. burnetii* effector CvpB perturbs PI 3-Kinase PIKFYVE activity, resulting in an accumulation of PI(3)P at CCVs(4). Indeed, 49% (±1.4%) of infected cells presented CCVs that were positive for PI(3)P at 2 days post-infection. This percentage increased to 74.4% (±1.5%) at 4 days post-infection, only to decrease slightly at 61.5% (±0.6%) at 6 days post-infection (Fig. 1A). At all time points of infections, over 90% of infected cells contained CCVs that were also positive for PI(4)P and LBPA (Fig. 1A, B). This latter is a typical marker of MVBs, where it serves an important role in the biogenesis of intraluminal vesicles (ILVs) by anchoring the ESCRT machinery protein ALIX to MVB membranes(14). All the other lipid sensors tested were largely absent from CCVs during the time course of infection (SI appendix, Fig. S1A).

**Figure 1:**
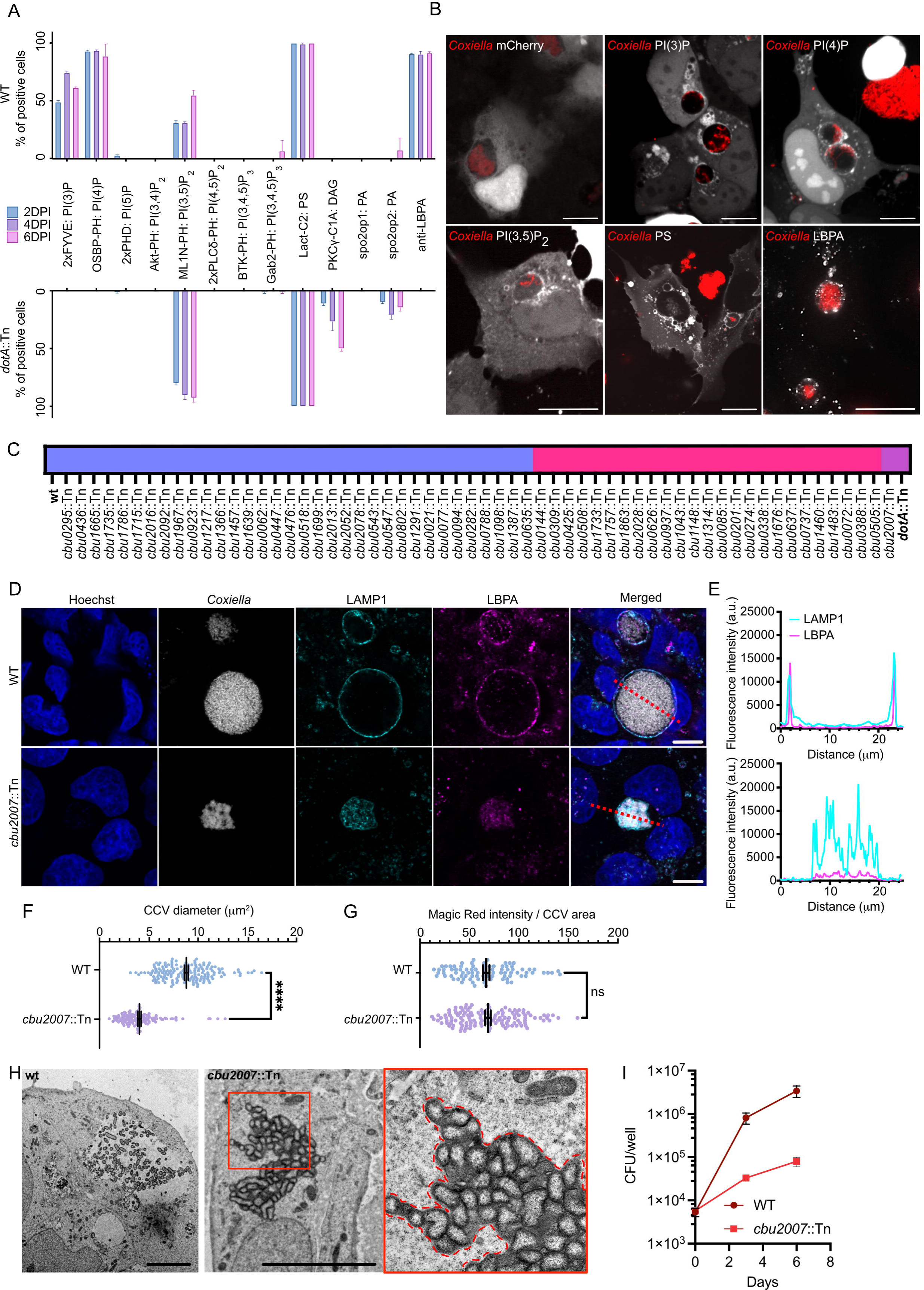
Lipid profiling of *Coxiella*-containing vacuoles identifies CBU2007 as an effector required for LBPA localization. (A) U2OS cells were infected with *Coxiella dotA*::Tn (bottom panel) or WT *Coxiella* (upper panel) for 2, 4 and 6 days and transfected with plasmids encoding a mCherry/RFP-tagged catalogue of lipid-binding sensors or labeled with an anti-LBPA antibody. The percentage of cells presenting CCVs positive for the indicated lipids was visually scored. Values are means ± standard deviation of duplicate experiments where 100 cells were scored for each condition, DPI: days post-infection. (B) Representative images of U2OS cells treated as described in (A) (*C. burnetii* is pseudocolored in red, lipid-binding sensors or LBPA labeling are pseudocolored in white). (C) U2OS cells were infected for 4 days either with a library of mutants carrying transposon insertions in validated or predicted effector-coding genes, with wt *C. burnetii* as positive control, or with the *dotA*::Tn mutant as a negative control, fixed and labeled with an anti-LBPA antibody. The presence of LBPA at the CCV was visually scored and ranked as compared to wt *C. burnetii* infected cells: Unaltered LBPA labeling of CCVs (blue), partial loss of LBPA labeling (pink) and complete loss of LBPA labeling (purple). (D) U2OS cells infected for 4 days either with wt *C. burnetii* (upper panel, grey) or the *cbu2007*::Tn mutant (bottom panel, grey) were labeled with Hoechst (blue), anti-LAMP1 (cyan) and anti-LBPA (magenta) antibodies. (E) The fluorescence intensity of LAMP1 (cyan) and LBPA (magenta) was measured along the red dash lines from (D). (F) The area of CCVs harboring either wt *C. burnetii* (WT) or the *cbu2007*::Tn mutant strain (*cbu2007*::Tn) was measured using LAMP1 labeling from U2OS cells infected for 4 days. Values are means ± standard deviation of three independent experiments (****P<0,0002; Unpaired t-test). (G) Proteolysis of cathepsin B-Magic Red was measured for individual CCVs at 4 days post-infection (dpi). Values are means ± standard deviation of three independent experiments (ns: non-significant; Unpaired t-test). (H) Representative electron micrographs of U2OS cells infected with wt *C. burnetii* (left panel) or the *cbu2007*::Tn mutant strain (right panel). The red inset shows an enlarged view of the CCV generated by the *cbu2007*::Tn mutant strain, dashed red line represents the CCV contour. (I) CFU counts from U2OS cells challenged with wt *C. burnetii* or the *C. burnetii cbu2007*::Tn mutant strain for 3 and 6 days. Scale bars: 10 µm

**Figure 2:**
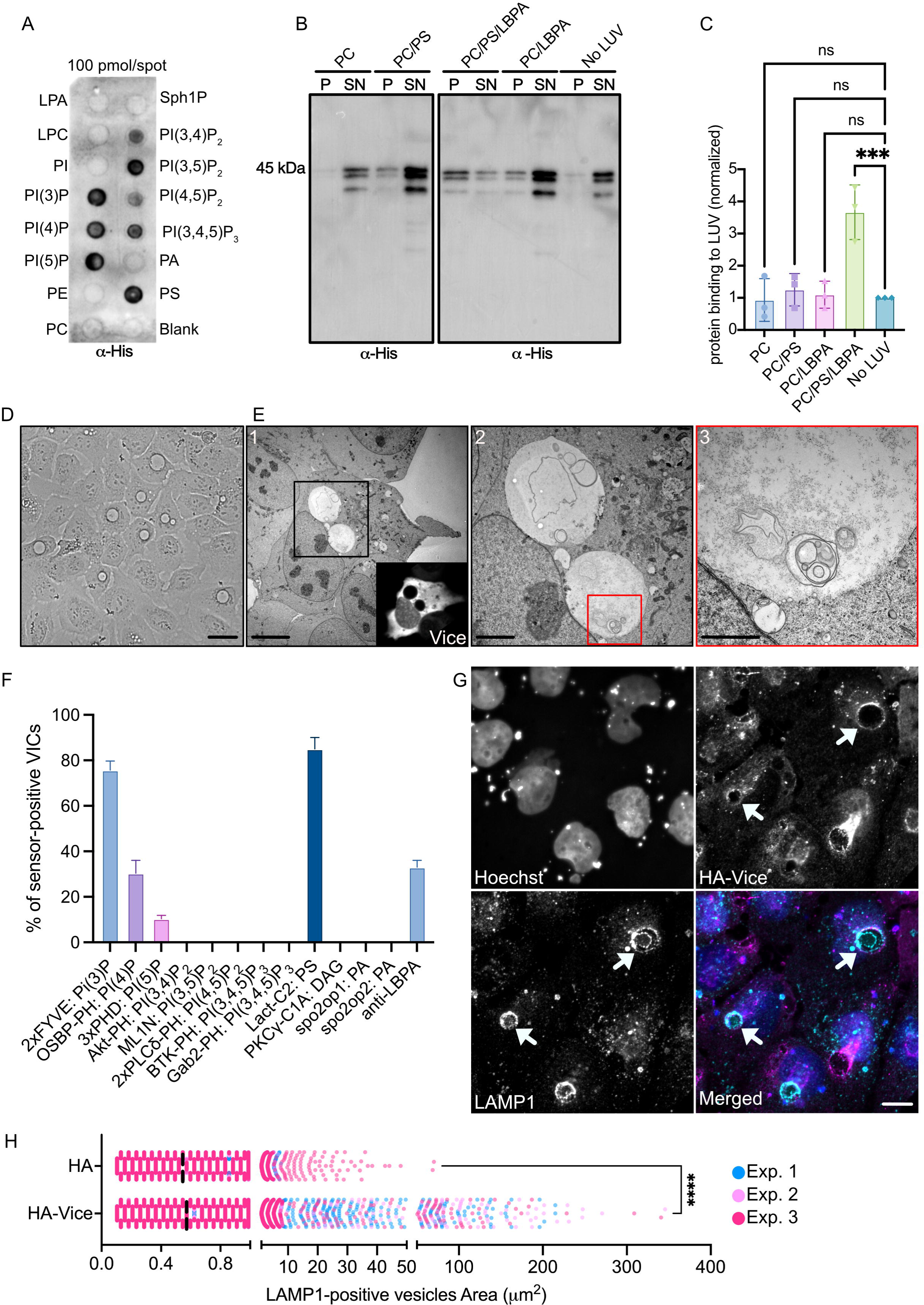
CBU2007/Vice is a lipid-binding effector involved in CCV biogenesis and LBPA enrichment. (A) Representative protein/lipid overlay assay performed with 6xHIS-CBU2007. (B) Representative Western blot of large unilamellar vesicles (LUVs) co-sedimentation assay using 6xHIS-CBU2007. CBU2007 in the pellet (P; bound to LUVs) or the supernatant (SN; unbound to LUVs) was detected using an anti-Histidine antibody. (C) CBU2007/LUVs binding was determined by measuring CBU2007 signal detected in (B) and using the following formula: (*P_intensity_ /* (*P_intensity_+ SN_intensity_*))*100. Values are means ± standard deviation of three independent experiments (***P<0,0001; ns: non-significant. One-way ANOVA, Dunnett’s multiple comparisons test). (D) Representative brightfield image of U2OS cells transfected for 24 hours with plasmids encoding HA-Vice, scale bar: 20 µm. (E) Correlative Light-Electron Microscopy (CLEM) of U2OS cells co-transfected with plasmids encoding HA-ALFA-Vice and NbALFA-mScarlet. Cells were imaged using confocal microscopy to identify the cell of interest (Vice inset) followed by fixation for electron microscopy ((1) scale bar: 5 µm; (2) scale bar: 2 µm; (3) scale bar: 1 µm). (F) The lipid composition of VICs was assessed by visually scoring the presence of the indicated lipid sensors in U2OS cells transfected with plasmids encoding HA-Vice and the indicated lipid sensors or HA-Vice alone followed by anti-LBPA labeling. Values are means ± standard deviation of three independent experiments where 100 VICs were scored for each condition. (G) U2OS cells transfected with a plasmid encoding HA-Vice (magenta) and labeled with anti-LAMP1 antibodies (cyan) indicate that VICs are positive for LAMP1 (white arrows). Hoechst was used to label nuclei (white) in all conditions. Scale bars: 10 µm. (H) The median size of LAMP1-positive compartments in U2OS cells expressing either HA-tag alone or HA-Vice was measured using CellProfiler. Each spot represents a LAMP1-positive compartment from three independent experiments (blue, pink, red; ****P<0,0001, unpaired t-test).

To determine whether *C. burnetii* effector proteins actively control the enrichment or exclusion of lipids at CCVs, we re-assessed the localization of all monitored lipids in cells challenged with the T4SS defective, *dotA*::Tn *C. burnetii* mutant for 2, 4 and 6 days. As expected, PS enrichment at vacuoles containing the *dotA*::Tn mutant remained unchanged as compared to CCVs harboring wt bacteria (Fig. 1A; SI appendix, Fig. S1B). Conversely, in agreement with our hypothesis, the percentage of PI(3,5)P_2_-positive CCVs was increased to 80.2 % (±1.3 %) at 2 days post-infection and further increased to over 90 % at later time points (Fig. 1A). Enrichment of PI(3,5)P_2_ correlated with a complete loss of PI(3)P at vacuoles containing the *dotA*::Tn mutant (Fig. 1A; SI appendix, Fig. S1B), as expected from our previous reports on the role of CvpB(4). Similarly, both PI(4)P and LBPA were completely absent from these compartments, indicating that their recruitment to CCV membranes is Dot/Icm-dependent (Fig. 1A). Finally, most of the lipids that were not observed at CCVs remained excluded from vacuoles harboring the *dotA*::Tn mutant, with the exception of diacylglycerol (DAG) and phosphatidic acid (PA), which are both present at the Golgi complex and the plasma membrane (Fig. 1A). Interestingly, PI(4)P (which we found enriched at CCVs but excluded from *dotA*::Tn-containing vacuoles) can be a source of DAG, through hydrolysis by phospholipase C (PLC)(15). In turn, DAG is the source of cellular PA. Thus, the loss of PI(4)P and the enrichment of DAG and PA at vacuoles containing the *dotA*::Tn mutants may be correlated. Together, our observations indicate that *C. burnetii* uses effector proteins to control the lipid composition of its replicative niche.

### Screening of C. burnetii effector proteins involved in the subversion of PI(4)P and LBPA trafficking

We previously reported that PI(3)P enrichment at CCVs depends on the translocation of the effector protein CvpB(4). Here, to identify *C. burnetii* effector proteins involved in PI(4)P and LBPA localization at CCVs, we challenged U2OS cells with a library of 60 GFP-tagged *C. burnetii* mutants carrying transposon insertions in genes encoding for candidate and validated effector proteins (Fig. 1C). PI(4)P was monitored by transfecting cells with a plasmid encoding mCherry-tagged OSBP-PH domain, whereas LBPA was monitored using anti-LBPA antibodies. Cells challenged with either wt *C. burnetii* or the *dotA*::Tn mutant (both expressing GFP) were used as positive and negative controls, respectively. At 4 days post-infection, mCherry-OSBP-PH-transfected cells were imaged by automated live confocal microscopy as described above, whereas non-transfected cells were fixed and labeled with anti-LBPA and anti-LAMP1 antibodies, prior to imaging. PI(4)P or LBPA localization was visually scored as described above to identify mutations that partially or completely impaired their accumulation at CCVs. None of the screened mutants showed a loss of PI(4)P enrichment at CCVs, suggesting that the manipulation of PI(4)P trafficking may be coordinated by effector proteins not included in our library. Conversely, 25 transposon insertions mildly affected LBPA accumulation at CCVs, and *C. burnetii* carrying a transposon insertion in *cbu2007* (*cbu2007*::Tn) completely failed to accumulate LBPA at CCVs (Fig. 1C, D, E; SI appendix, Fig. S2). Based on LAMP1 labeling, wt CCVs presented a diameter spanning from 5 to 15 μm, while CCVs generated by the *cbu2007*::Tn mutant were 1 to 8 μm in size, revealing a significant reduction in CCVs area (Fig. 1F). The proteolytic activity of CCVs harboring the *cbu2007*::Tn mutant, which is directly correlated with CCV acidification, was indistinguishable from that of CCVs harboring wt *C. burnetii* (Fig. 1G), indicating proper CCV metabolic activation and T4SS secretion of the *cbu2007*::Tn mutant. Electron microscopy revealed that U2OS cells challenged with the *cbu2007*::Tn mutant strain were pleomorphic, with membranes tightly sealed against compact bacterial colonies, while the wt bacterium developed large, spacious CCVs, suggesting a role for CBU2007 in CCV biogenesis (Fig. 1H). We were unable to complement the *cbu2007* mutation by inserting a wt copy of the gene in the mutant strain. Regardless, ectopic expression of HA-tagged CBU2007 in cells challenged with the *cbu2007*::Tn mutant strain restored the shape of CCVs and the localization of LBPA (SI appendix, Fig. S3A, B). We also transformed *Legionella pneumophila* with plasmids encoding 1-Lactamase-tagged Vice (BLAM-Vice) to assess effector translocation by a BLAM assay and to determine whether Vice alone was sufficient to trigger the expansion of other intracellular bacteria niches. Vice was indeed translocated by the *Legionella* T4SS (SI appendix Fig. 3C) and bacteria expressing BLAM-Vice replicated in larger vacuoles (SI appendix Fig. 3D). The impact of the CCV biogenesis defect associated with the mutation of *cbu2007* on infections was assessed by colony-forming units (CFU) counts from U2OS cells challenged with either wt *C. burnetii* or the *cbu2007*::Tn mutant strain for 3 and 6 days. The mutation in *cbu2007* dramatically reduced *C. burnetii* intracellular replication at both time points of infection (Fig. 1I). Collectively, these observations indicate that CBU2007 is important for infection and involved in the initial expansion of CCVs.

**Figure 3:**
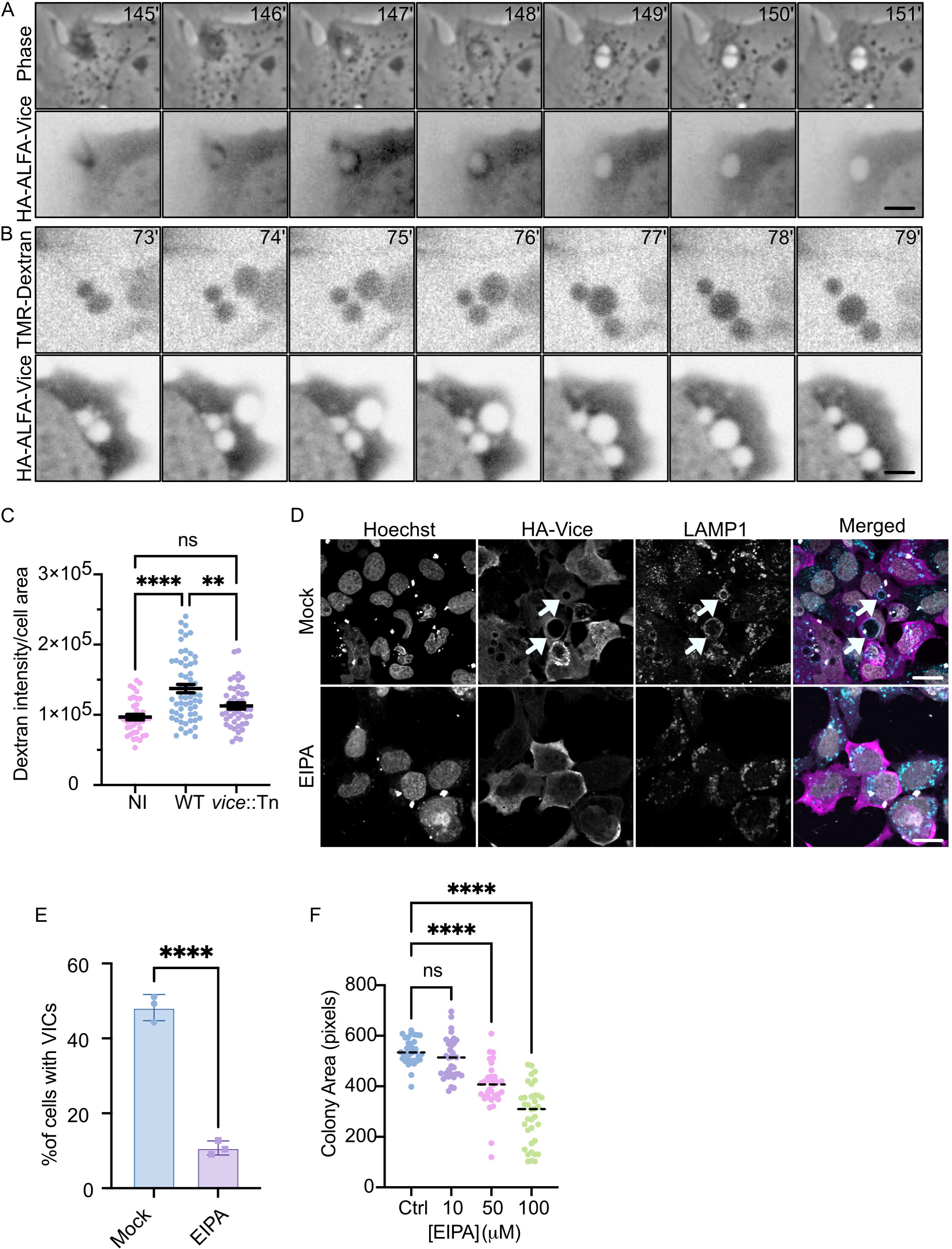
Vice stimulates macropinocytosis. (A) Representative images from Movie S1 illustrating U2OS cells co-transfected with plasmids encoding HA-ALFA-Vice and NbALFA-mScarlet and imaged after 8 hours in phase contrast (upper panels) and 555 nm emission (lower panels) channels. Scale bars: 4 µm. (B) Representative images from Movie S6 illustrating U2OS cells co-transfected with plasmids encoding HA-ALFA-Vice and NbALFA-mNeonGreen for 8 hours prior to TMR-Dextran addition. Images were acquired every minute overnight in 555 nm emission (upper panel) and 488 nm emission (lower panel) channels. Scale bars: 4 µm. (C) Stimulation of macropinocytosis was assessed by measuring the fluorescence intensity of TMR-Dextran from non-infected (NI), wt-infected (WT) or vice::Tn mutant-infected (*vice*::Tn) U2OS cells. Values are means ± standard deviation of three independent experiments (****P<0,0001, **P<0,0022, ns: non-significant. One-way ANOVA, Dunnett’s multiple comparisons test). (D) U2OS cells transfected with a plasmid encoding HA-Vice were incubated overnight with either methanol (Mock, upper panel) or EIPA (lower panel). VIC formation was assessed by using Hoechst to label nuclei (white), anti-HA (magenta) and anti-LAMP1 antibodies (cyan). White arrows point at VICs; scale bars: 20 µm. (E) The percentage of cells described in (D) containing VICs was visually scored. Values are means ± standard deviation of three independent experiments (****P<0,0001, unpaired t-test). (F) U2OS infected with wt *C. burnetii* were incubated with the indicated concentration of EIPA for 48 hours. The median area of *C. burnetii* colonies was measured from 25 images per condition (corresponding to more than 3000 colonies per condition). Values are means of two independent experiments (****P<0,0001, ns: non-significant. One-way ANOVA, Dunnett’s multiple comparisons test).

### CBU2007 is a T4SS C. burnetii effector important for CCVs expansion

*cbu2007* encodes the *C. burnetii* protein with the highest effector prediction score assigned by the S4TE, an algorithm for T4SS effectors prediction(16). Its expression is regulated by a PmrA promoter and it includes a putative C-terminal E-Block secretion signal(17). CBU2007 has been recently validated as a secreted *C. burnetii* effector protein conserved among *C. burnetii* genomes(18), is important for CCV biogenesis(18) and essential for *C. burnetii* survival in primary macrophages(19). Nested PCR followed by sequencing(20) indicated that the transposon insertion occurred 711 bp downstream of the gene starting codon. Southern blot analysis validated the presence of a single transposon insertion in the genome of the *cbu2007*::Tn strain (SI appendix, Fig. S3E). To confirm CBU2007 translocation, *cbu2007* was cloned into the pXDC61K-BLAM vector to generate an N-terminal fusion with β-lactamase, and transformed either into GFP-tagged *C. burnetii* (wt) or the T4SS-defective *dotA*::Tn mutant strain. The expression of β-lactamase-tagged constructs was validated by Western blot using an anti-β-lactamase antibody (SI appendix, Fig. S3F). Protein translocation in U2OS cells was then assessed at 24, 48 and 72 h post-infection by means of a β-lactamase secretion assay. *C. burnetii* expressing β-lactamase alone or β-lactamase-tagged CvpB(4) were used as negative and positive controls, respectively. CBU2007 was efficiently translocated at all time points of infection, whereas no translocation was observed in cells challenged with the T4SS-defective mutant strain (SI appendix, Fig. S3G), indicating that CBU2007 is indeed a T4SS substrate.

### CBU2007 interacts with host lipids

As CBU2007 is required to accumulate LBPA at CCVs, we generated Histidine-tagged recombinant CBU2007 to test the lipid-binding properties of the *C. burnetii* effector protein. First, PIP Strips were overlaid with 6xHis-CBU2007 and labeled with an anti-His antibody. This revealed an interaction of 6xHis-CBU2007 with PS and phosphorylated PIs, with a preferential binding to PI(3)P, PI(4)P and PI(5)P (Fig. 2A). No interactions were observed when PIP Strips were incubated with purified 6xHis-GST (SI appendix, Fig. S3H). As commercially available PIP Strips do not include LBPA, an *in vitro* co-sedimentation assay using Large Unilamellar Vesicles (LUVs) was used to assess the binding of CBU2007 to biological membranes containing LBPA. Co-sedimentation of 6xHis-CBU2007 with LUVs containing 80% phosphatidylcholine (PC) in combination with either 20% LBPA or 20% PS (Fig. 2B, C) was not significant. Conversely, 6xHis-CBU2007 efficiently co-sedimented with vesicles containing a mixture of 70% PC, 10% PS and 20% LBPA (Fig. 2B, C), which is compatible with the lipid composition of late endosomes and MVBs. In the absence of LUVs, His-CBU2007 was only detected in the supernatant, indicating that the protein does not precipitate (Fig. 2B, C). Together, these data indicate that CBU2007 can interact with PS and LBPA-enriched membranes.

### CBU2007 is a vacuole-inducing effector protein

To assess the role of CBU2007 in manipulating host cell functions leading to CCV expansion, we ectopically expressed the effector protein in U2OS cells. Remarkably, the vast majority of cells expressing CBU2007 presented large, circular vacuoles, whose morphology was reminiscent of early CCVs observed in *C. burnetii*-infected cells (Fig. 2D). VIC formation was observed in multiple cell types ectopically expressing Vice but was more prominent in U2OS and A549 cells (SI appendix, Fig. S4A). Correlative electron microscopy revealed that these compartments are spacious, single-membrane organelles, containing luminal vesicles of a size ranging from 200 nm to 2 µm (Fig. 2E). Organelles reminiscent of lamellar bodies were also observed in the lumen of the compartments (Fig. 2E). Given this remarkable phenotype, we renamed the *C. burnetii* effector Vice (for Vacuole-Inducing *Coxiella* Effector). Co-expression of HA-Vice with lipid-binding sensors, or immunolabeling of HA-Vice-expressing cells with anti-LBPA antibodies, revealed that the membranes of Vice-induced compartments (VICs) contain PS, LBPA, and the monophosphorylated PIs interacting with Vice indicated above (PI(3)P, PI(4)P and PI(5)P; (Fig. 2F). VICs are also positive for LAMP1 (Fig. 2G), but negative for Lysotracker (SI appendix, Fig. S4B), indicating that they may derive from late endosomes. HA-Vice itself was observed decorating VIC membranes in over 50% of transfected cells (Fig. 2G). Finally, LAMP1 labeling was used to determine the size of late endosomes and lysosomes in U2OS cells ectopically expressing HA-Vice or the HA tag alone as a control. A population of LAMP1-positive compartments with a median area of 0.5 µm^2^ (corresponding to a diameter of ∼0.8 µm) was observed in both conditions (Fig. 2H). In addition, cells overexpressing Vice also presented a population of much larger compartments with an area ranging from 50 to 250 µm^2^ (corresponding to a diameter of 8-17 µm; Fig. 2H). We thus defined VICs as intracellular compartments with a diameter ≥ 8 µm.

### Vice enhances macropinocytosis

To monitor the biogenesis of Vice-induced compartments (VICs) by live imaging, we generated a plasmid encoding ALFA-tagged Vice, allowing the imaging of the *Coxiella* effector in cells expressing fluorophores-coupled ALFA-targeting nanobodies(21). Cells were imaged either at early time points after transfection (8h), to observe the initial formation of VICs, or at later time points (24h), to monitor their dynamics and persistence in cells. Early after transfection, HA-ALFA-Vice was mostly cytoplasmic and few large compartments were observed in the cytoplasm of transfected cells (SI appendix, Fig. S4C), while ectopic expression of the anti-ALFA nanobodies alone never resulted in VICs formation (SI appendix, Fig. S4C). Surprisingly, HA-ALFA-Vice transiently accumulated at defined regions of the plasma membrane, which was systematically accompanied by the formation of large vacuoles (Movie S1; Fig. 3A). These remained labeled by HA-ALFA-Vice for a short period of time (1-3 min), after which HA-ALFA-Vice rapidly redistributed to the cytoplasm (Movie S1; Fig. 3A) and the newly formed compartments moved to a perinuclear region, where they sometimes fused with other pre-existing compartments. Vice accumulation at the plasma membrane and the internalization of large vacuoles were never observed at later time points of transfection. Instead, live cell imaging revealed a remarkable stability of VICs, which persisted unperturbed in cells over 8-10 hours (Movie S2).

Given the remarkable size of VICs, we hypothesized that Vice could enhance macropinocytosis. HA-ALFA-Vice and anti-ALFA-mScarlet nanobodies were thus ectopically expressed in U2OS cells in combination with either the PI(4,5)P_2_ sensor 2xPLCδ-GFP, GFP-LifeAct to visualize filamentous actin (F-actin), or GFP-RAB7. Vice accumulation at the plasma membrane was slightly preceded by a strong accumulation of PI(4,5)P_2_, which persisted at VICs for a few minutes after the cytoplasmic redistribution of Vice (Movie S3; SI appendix, Fig. S5A). Conversely, Vice and actin accumulated simultaneously at the plasma membrane (Movie S4; SI appendix, Fig. S5B). However, actin persisted at VICs long after Vice dissociation (Movie S4; SI appendix, Fig. S5B). Finally, GFP-RAB7 was observed at VICs rapidly after the closure of the plasma membrane and remained associated with VICs after the dispersion of Vice to the cytoplasm (Movie S5; SI appendix, Fig. S5C). Next, U2OS cells ectopically expressing HA-ALFA-Vice and anti-ALFA-mNeonGreen nanobodies were incubated overnight with 0.5 mg/ml tetramethylrhodamine (TMR)-tagged 70kDa dextran, washed in PBS and imaged over 8 hours to monitor fluid phase macropinocytosis. Large TMR-dextran-positive macropinosomes formed in Vice-expressing cells and often fused together to form larger VICs which persisted in cells over the time course of the experiment (Movie S6; Fig. 3B). Importantly, U2OS cells infected with wt *Coxiella* incorporated significantly higher levels of 70kDa dextran as compared to non infected or *vice*::Tn-infected cells, indicating that Vice stimulates macropinocytosis during infection (Fig. 3C). Finally, U2OS cells ectopically expressing HA-ALFA-Vice and anti-ALFA-mScarlet nanobodies were incubated overnight with 50 µM of 5L(*N*LethylL*N*Lisopropyl)amiloride (EIPA), which specifically inhibits macropinocytosis(22). In agreement with our hypothesis, EIPA treatment dramatically reduced the formation of VICs (Fig. 3D, E) and triggered an accumulation of Vice at the plasma membrane (Fig. 3D). Accordingly, EIPA significantly affected the biogenesis of CCVs in cells infected with wt *Coxiella* and incubated with increasing concentrations of the macropinocytosis inhibitor 24 hours post infection (Fig. 3F; SI appendix Fig. S5D). Our data indicate that Vice is transiently recruited at the plasma membrane of eukaryotic cells where it triggers macropinocytosis, leading to the formation of large intracellular compartments whose morphology and composition are reminiscent of early CCVs. In the context of infections, this may provide CCVs with a source of membrane required for the initial rapid expansion of the *C. burnetii* replicative niche.

### Vice perturbs the ESCRT machinery

As described above, VICs can persist within cells over several hours without significantly altering their morphology and protein/lipid composition. Given that VIC membranes contain LBPA (Fig. 2F) and RAB7 (SI appendix, Fig. S6A), typical markers of multivesicular bodies (MVBs)(14), we hypothesized that Vice may also interfere with the ESCRT machinery to inhibit membrane invagination and pinching off of ILVs to stabilize VICs. We thus investigated the intracellular localization of GFP-ALIX and the ESCRT-I component TSG101 in U2OS cells ectopically expressing either HA-Vice or the HA tag alone as a negative control. Both GFP-ALIX and TSG101 were observed in spots in the cytoplasm of HA-expressing cells, suggesting membrane recruitment at MVBs (Fig, 4A, B, SI appendix, Fig. S6B, C). TSG101 decorated the contour of approximately 50% of VICs in cells expressing HA-Vice (SI appendix, Fig. S6B), whereas GFP-ALIX was mainly cytoplasmic when co-expressed in combination with HA-ALFA-Vice (Fig. 4A, B), suggesting that Vice may interfere with ALIX membrane anchoring, thereby perturbing the pinching off of ILVs.

**Figure 4:**
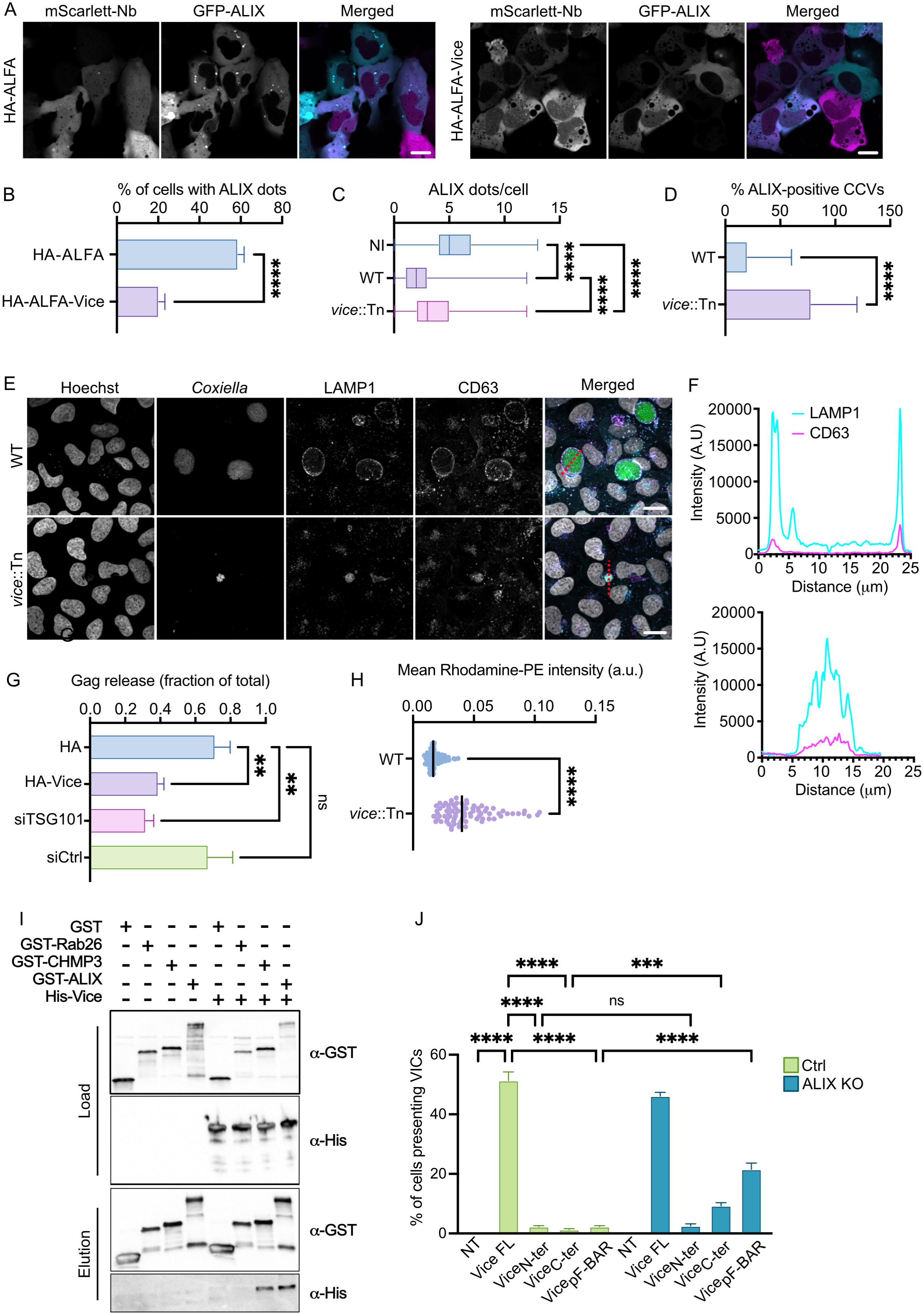
Vice inhibits the activity of the ESCRT machinery. (A) U2OS cells were co-transfected with plasmids encoding GFP-ALIX (cyan), NbALFA-mScarlet (magenta) and either HA-ALFA (left panel)s or HA-ALFA-Vice (right panels). (B) The percentage of cells presenting GFP-ALIX dots in cells treated as in (A), was visually scored. (C) U2OS cells infected for 4 days with either WT *Coxiella* or *vice*::Tn *C. burnetii* mutant were transfected for 24h with a plasmid encoding GFP-ALIX and the number of GFP-ALIX dots in non-infected (NI) and infected cells was visually scored. (D) GFP-ALIX localization at CCVs was visually scored in both infection conditions described in (C). Values are means ± standard deviation of three independent experiments (****P<0,0001, unpaired t-test for B and D, One-way ANOVA, Dunnett’s multiple comparisons test for C). (E) U2OS cells infected for 4 days with either WT *Coxiella* or *vice*::Tn *C. burnetii* mutant (green) were labeled with Hoechst (white), anti-LAMP1 (cyan) and anti-CD63 (magenta). (F) Representative distribution of LAMP1 (cyan) and CD63 (magenta) fluorescence intensity was measured along the corresponding red dash lines illustrated in (F). Scale bars: 20 µm. (G) HEK293T were co-transfected with plasmids encoding either HA or HA-Vice and GAG, or transfected with plasmids encoding GAG, 24 hours after transfection with either TSG101-targeting siRNAs (siTSG101) or scrambled siRNA (siCTRL). Virus-like particles (VLP) release was assessed by Western blot using anti-p24 antibodies and the relative percentage of VLPs was measured by quantifying the p24 signal from triplicates experiments using the following formula: (VLPs*_p24_intensity_ /* (VLPs*_p24_intensity_ +* Cells*_p24_intensity_*)). Values are means ± standard deviation of three independent experiments (**P<0,001, ns: non-significant, One-way ANOVA, Dunnett’s multiple comparisons test). (H) The CCV-associated fluorescence intensity of Rhodamine-PE was measured from maximum intensity projection of confocal images of U2OS cells infected either with WT *C. burnetii* (WT) or the *vice*::Tn mutant strain (*vice*::Tn). Values are mean of three independent experiments where 100 CCVs were measured for each condition (****P<0,0001, Unpaired t-test). (I) Protein co-purification on agarose-glutathione resin from lysates of LEMO21 bacteria transformed with pGEX4T1 (GST), pGEX4T1-RAB26 (GST-RAB26), pGEX4T1-CHMP3B (GST-CHMP3) or pGEX4T1-ALIX (GST-ALIX) in combination with pTWIST-Vice (His-Vice). Total cell lysates (Load) and purified proteins (Elution) were labeled with anti-GST and anti-His antibodies. (J) Occurrence of VICs was visually scored from control (Ctrl) or ALIX-KO (ALIX KO) U2OS cells transiently expressing the indicated HA-tagged Vice truncations or the HA tag alone as control. Values are means ± standard deviation of three independent experiments (****P<0,0001, ***P<0,0022, ns: non-significant, One-way ANOVA, Dunnett’s multiple comparisons test).

Similar phenotypes were recapitulated in U2OS cells infected with wt *C. burnetii*. Infections of U2OS cells by either the wt strain or the *vice*::Tn mutant significantly decreased the number of ALIX dots in the cytoplasm, with a more substantial effect observed in cells infected by the wt strain (Fig. 4C). More importantly, when cells were exposed to the *vice*::Tn mutant strain, 78% of CCVs were decorated by ALIX, as opposed to 20% of CCVs in cells challenged with the wt strain (Fig. 4D, SI appendix Fig. S6D). Finally, we investigated the intracellular localization of CD63, a tetraspanin typically found at MVBs and ILVs(23). CD63 was observed in spots in the cytoplasm of HA-expressing U2OS cells and decorated the contours of VICs in HA-Vice-expressing cells (SI appendix, Fig. S7A). CD63 was also observed within the lumen of VICs (SI appendix, Fig. S7A), in agreement with our CLEM observations reported in Fig. 2E. CD63 decorated the contour of CCVs generated by wt *C. burnetii*, while the protein was observed within the lumen of CCVs generated by the *vice*::Tn mutant strain (Fig. 4E, F). Ectopic expression of Vice in U2OS cells challenged with the *vice*::Tn mutant strain restored CCV morphology and CD63 localization (SI appendix, Fig S7B, C). It has been previously reported that the ESCRT machinery also regulates HIV release from infected cells(24). Thus, to further investigate the possibility that Vice alters the functionality of the ESCRT machinery, we ectopically expressed the HIV polyprotein Gag in HEK293T cells, in combination with either the HA tag alone or HA-Vice, and monitored the levels of extracellular Gag as a proxy for the release of pseudo-viral particles. HEK293T cells transfected with Gag and siRNA targeting TSG101, an essential component of the ESCRT machinery previously shown to inhibit the release of viral particles(25), were used as control. In agreement with our hypothesis, cells depleted of TSG101 or overexpressing HA-Vice released significantly less Gag in the supernatant as compared to cells expressing Gag in combination with either non-targeting siRNAs or the HA tag alone (Fig. 4G, SI appendix Fig. S7D). To further assess the inhibition of the ESCRT machinery we monitored the incorporation of Rhodamine-PE by cells (as a proxy for ILVs formation(26)) in the context of ectopic expression of HA-Vice or infection with either wt *Coxiella* or the *vice*::Tn mutant strain. Confocal microscopy revealed that Rhodamine-PE labeled the contour of VICs but was not observed within their lumen (SI appendix Fig. S8A). Accordingly, CCVs harboring the *vice*::Tn mutant strain were Rhodamine-PE-positive, indicating the presence of ILVs, as opposed to CCVs generated by wt bacteria, which were largely Rhodamine-PE negative (Fig. 4H, SI appendix Fig. S8B). In addition, yeast two-hybrid screening identified the ESCRTIII component CHMP3 as a candidate interactor of Vice. Thus, we aimed at validating interactions of Vice with ESCRT components by protein co-purification in *E. coli*. Of note, we could show that Vice interacts with CHMP3 and ALIX, whereas no interactions were observed between Vice and GST alone or GST-RAB26 used as negative controls (Fig. 4I).

### Functional prediction of Vice domains

Given the multifunctional role of Vice we carried out a bioinformatics analysis of its amino acid sequence to identify putative eukaryotic-like domains. The RaptorX algorithm(27) identified a putative F-BAR domain spanning aa 214-345 (SI appendix Fig. 8C). We have thus generated HA- and ALFA-tagged truncation mutants of Vice corresponding to the N-terminal domain (aa 1-213; Vice_N-ter_), the C-terminal domain (aa 214-395; Vice_C-ter_) and the predicted F-BAR domain (aa 214-345; Vice_pF-BAR_), with the aim of ascribing specific functions to different protein domains. None of the mutants led to the appearance of VICs when ectopically expressed in cells (Fig. 4J). Vice_C-ter_ and Vice_pF-BAR_ mostly accumulated at the plasma membrane whereas Vice_N-ter_ was diffused in the cytoplasm (SI appendix Fig. 9). Live imaging of cells expressing the ALFA-tagged truncation mutants in combination with the PI(4,5)P_2_ probe 2XPLCδ-PH domain confirmed the plasma membrane accumulation of Vice_C-ter_ and the Vice_pF-BAR_ and revealed that both mutants induced the internalization of large compartments positive for both PI(4,5)P_2_ and the effector mutants (Movie S7 and S8). These compartments were not stable and rapidly shrank upon internalization, suggesting that the C-terminal domain of Vice may stimulate macropinocytosis. Conversely, conventional-sized endosomes internalized by cells ectopically expressing Vice_N-ter_ and the PI(4,5)P_2_ probe were occasionally growing in size following internalization, suggesting that the N-terminal domain of Vice may stabilize endosomal membranes (Movie S9). To validate our hypothesis, we generated CRISPR-Cas9 edited U2OS cells to suppress the expression of endogenous ALIX (SI appendix Fig. 10A). ALIX KO did not affect the formation or the morphology of VICs in cells expressing full length Vice, nor restored the formation of VICs in cells expressing Vice_N-ter_ (Fig. 4J, SI appendix Fig. 9). Conversely, ectopic expression of either Vice_C-ter_ or Vice_pF-BAR_ partially restored VICs formation in ALIX-KO cells (Fig. 4J, SI appendix Fig. 9). When tested in the context of infection, ALIX-KO did not affect the size or morphology of CCVs harboring wt bacteria, nor rescued the CCVs morphology phenotype observed with *vice*::Tn mutant strain (SI appendix Fig. 10B), suggesting indeed that the double function of the effector protein (stimulation of macropinocytosis and inhibition of the ESCRT machinery) are important for CCVs biogenesis.

## Discussion

Given the importance of PI(3)P in CCV biogenesis and in *C. burnetii* virulence *in vivo*, in this study, we used lipid-binding fluorescent sensors and antibodies to further characterize CCV lipid composition. PI(3,5)P_2_ and PS, which are typically found at late endosomes/lysosomes, were observed at CCVs harboring wt bacteria or the T4SS-defective mutant *dotA::*Tn, suggesting that *C. burnetii* does not alter their trafficking. On the contrary, the late-endosome/MVB lipid marker LBPA, the early endosomal lipid PI(3)P and the Golgi-associated lipid PI(4)P were specifically enriched at CCVs generated by wt bacteria but absent from vacuoles containing the *dotA*::Tn mutant, indicating that *C. burnetii* actively manipulates their trafficking/metabolism. Cross-referencing the data obtained with lipid sensors we could also observe that vacuoles harboring the *dotA*::Tn mutant were negative for PI(3)P as previously reported(4), but enriched in PI(3,5)P_2_, supporting our hypothesis that CvpB interferes with PI(3)P phosphorylation to PI(3,5)P_2_. Phenotypic screen of a library of *C. burnetii* mutants carrying transposon insertions in genes encoding predicted and validated effector proteins allowed us to pinpoint a transposon insertion in *cbu2007*, which largely inhibited LBPA localization at CCVs. CBU2007 is the highest-scoring predicted effector by the S4TE algorithm(16) and has been recently validated as a *C. burnetii* effector protein(18). CBU2007 is required for intracellular replication(18) and, more importantly, is essential for survival in primary macrophages and virulence in mice(19).

Membranes of CCVs harboring *cbu2007::Tn* mutants are collapsed on bacteria as opposed to the spacious CCVs generated by wt *C. burnetii*. Despite their altered morphology, collapsed CCVs are positive for the late endosomal/lysosomal marker LAMP1 and are acidified, similar to wt CCVs, suggesting that bacterial metabolism and effector translocation are not affected. Indeed, differently from *dotA*::Tn mutants, the *cbu2007*::Tn mutant strain replicates within these compartments, albeit much less than wt bacteria.

Most strikingly, ectopic expression of CBU2007 in epithelial cells leads to the formation of a very large vacuole, with a diameter ranging from 8 to 17 µm, resembling early CCVs in both morphology and protein/lipid composition. We thus named the effector encoded by *cbu2007* Vice (for <u>V</u>acuole-inducing *<u>C</u>oxiella* <u>e</u>ffector). Recombinant Vice interacts with phosphoinositides and PS, indicating a possible association with the plasma membrane and late endosomal compartments. Vice co-sedimentation with LUVs enriched in PS and LBPA is suggestive of an interaction with MVBs. Combined with the observation of LBPA at CCVs generated by wt bacteria and the absence of LBPA from CCVs generated by the *cbu2007*::Tn mutant, our data indicate an interaction between MVBs and CCVs.

Given their morphology and lipid composition, we initially believed that Vice-induced compartments (VICs) could derive from MVBs failing to pinch off intraluminal vesicles, thereby increasing their membrane surface. We thus imaged cells ectopically expressing Vice over time to explore the biogenesis of VICs formation. To our surprise, VICs are not formed by the expansion of intracellular vesicles, but rather originate from the plasma membrane by macropinocytosis-like events. Live imaging also highlighted a dynamic association of Vice with membranes, the effector protein being mainly cytoplasmic, with transient accumulation at the plasma membrane at VICs internalization sites, followed by a dissociation from VICs shortly after internalization (1-2 min). The presence of macropinocytic markers and the inhibition of VIC formation by the macropinocytosis inhibitor EIPA, suggest that Vice stimulates macropinocytosis in cells. As recruitment of Vice at the plasma membrane follows the local accumulation of PI(4,5)P_2_ at the macropinocytic cup, we hypothesized that Vice recruitment may depend on interactions with specific lipids or proteins. Based on our observations on the ectopic expression of Vice, here we also show for the first time that macropinocytosis is essential for *C. burnetii* infections as EIPA effectively inhibits CCVs biogenesis.

Following internalization, VICs move to a perinuclear area where they remain remarkably stable in size and shape over several hours, suggesting that perturbing ILVs formation, which we envisioned as a biogenesis strategy, may in fact be involved in stabilizing these structures. Indeed, besides RAB7, CD63 and LBPA which are typically found at late endosomes and MVBs (28, 29), components of the ESCRT machinery, which normally cycle between late endosomal membranes and the cytoplasm, accumulate at the periphery of VICs, suggesting a perturbation of the machinery itself. Indeed, the typical ILV/exosomal marker CD63 decorates the contours of CCVs and VICs, whereas it is found within the CCV lumen in cells infected by the *vice*::Tn mutant strain. Membrane association of the ESCRT-associated protein ALIX, which is involved in pinching off ILVs from MVBs membranes, seems largely impaired in cells expressing Vice. Similar phenotypes, albeit attenuated, are also observed in the context of infection. Only 20% of CCVs generated by the wt strain are positive for ALIX whereas the majority of CCVs harboring the *vice::*Tn mutant strain are also positive for ALIX, suggesting an active ESCRT machinery pinching off vesicles from the CCV harboring the mutants strain, possibly leading to the convoluted, collapsed morphology observed. Accordingly, CCVs from cells infected with the *vice*::Tn mutant strain are positive for Rhodamine-PE, indicating the presence of ILVs, whereas PE labeling is limited to the contours of CCVs hosting wt bacteria. The milder phenotype observed during infection is probably related to the lower amounts of endogenous Vice translocated by bacteria as opposed to the ectopic expression of Vice in non-infected cells.

In support of a perturbation of the ESCRT machinery, ectopically expressing Vice significantly reduces the release of Gag-containing Viral-like particles, which is also mediated by the activity of the ESCRT machinery at the plasma membrane(24). The precise molecular mechanisms mediating the inhibition of the ESCRT machinery by Vice remains to be defined. However, we report here that Vice interacts with LBPA, ALIX and CHMP3, suggesting a direct interference with the assembly of the ESCRT complex.

The recent implication of the ESCRT machinery in the repair of CCV membranes during infections may seem counterintuitive to our findings(30). On the contrary, it has also been reported that in the context of membrane repair, membrane recruitment of ALIX solely relies on ALG-2 and LBPA is never observed at sites of membrane damage(31, 32). Therefore, multiple ESCRT-dependent machineries are involved in membrane repair and MVBs biogenesis, and perturbating the LBPA-dependent pathway allows *C. burnetii* to selectively target the pathway related to ILV biogenesis. Our hypothesis is that Vice association with membranes enriched in PS and LBPA might interfere with ALIX association with membranes, which is similarly mediated by binding to LBPA. During MVB biogenesis, LBPA is also found at ILV membranes within the lumen of MVBs. The perturbation of ILV formation in cells exposed to Vice, either by ectopic expression or by translocation during infections, would lead to the observed accumulation of LBPA at the surface of VICs or CCVs, respectively. This work also adds to our knowledge of the role of MVBs and the ESCRT machinery in infections by intracellular pathogens. Indeed, it has been previously reported that *M. tuberculosis* needs to perturbs the ESCRT machinery notably to escape into the cytoplasm(33, 34). It has also been reported that macrophages infected with *M. avium* or *M. smegmatis* secrete more exosomes as compared to non-infected cells(35). Secreted exosomes contain more Hsp70, which triggers the NF-LB signaling pathway in bystander cells(35). Thus, inhibiting the ESCRT machinery may also be a strategy for stealth pathogens to dampen inflammation. On the other hand, *Anaplasma phagocytophilum-*containing vacuoles (ApVs) are positive for MVB markers including LBPA and CD63, similar to what we have observed here(36). Indeed, MVB biogenesis is important for *A. phagocytophilum* replication within infected cells. Contrary to *C. burnetii*, however, *A. phagocytophilum* uses exosomes to release infectious progeny to bystander cells(36). Accordingly, ApV membranes are positive for ALIX, suggesting active ESCRT machinery, as opposed to what is observed here with CCVs(36).

In conclusion, here we report how lipid profiling of CCVs led to the identification and characterization of a new multi-functional *C. burnetii* effector protein Vice (CBU2007). Intracellular bacterial pathogens secreting effector proteins with multiple functions during infections have previously been reported, with *Chlamydia* being one of the best examples(37). Regarding *C. burnetii*, the effector protein CaeB was yet the only multi-functional effector described(38–41).

Here we propose that early during infection Vice promotes macropinocytosis to deliver membranes and nutrients to support the rapid expansion of early CCVs. At later time points of infection, Vice perturbs the ESCRT machinery by hindering membrane recruitment of ALIX, thereby limiting the formation of ILVs and stabilizing CCVs morphology. Additional work will focus on the precise molecular mechanism regulating Vice functions and will determine whether Vice may be involved in decreasing the release or altering the composition of extracellular vesicles by cells infected with *C. burnetii*.

## Acknowledgements

The authors are grateful to: Raphael Gaudin, Maïka Deffieu (IRIM CNRS UMR9004, Montpellier, France), Cécile Gauthier-Rouvière (CRBM, CNRS UMR5237), Anne Bonhoure and Michel Vidal (LPHI, CNRS UMR5294) for providing plasmids and for scientific discussions. We acknowledge the imaging facility MRI, member of the national infrastructure France-BioImaging supported by the French National Research Agency (ANR-10-INBS-04, «Investments for the future»). This work was supported by the French National Research Agency (ANR; ANR-17-CE15-0021, project QPiD).

