## Supplemental information for "The multifunction *Coxiella* effector Vice stimulates macropinocytosis and interferes with the ESCRT machinery"

This PDF file includes:

Supplementary text

Supplementary Figures S1 to S10

Supplementary Movies Legend S1 to S9

Supplementary Tables S1 to S4

**Supplementary Information Text**

**Extended Methods**

*Antibodies and reagents*

Mouse antibodies used were against Beta-Lactamase (BLAM) (1:2000, MA120370, ThermoFisher Scientific), GAPDH (1:1000, G8795, Sigma), HA (1:1000, 66006-1-Ig, Proteintech), CD63 (WB) (1:50, clone R5G2, MBL), CD63 (IF) (1:200 clone CLB-gran/12, 435, Sanquin), TSG101 (IF) (67381-1-Ig, Proteintech), GST-HRP (1:2000, 16-209, Sigma), HIS-HRP (1:2000, HIS.H8, ThermoFisher Scientific). Rabbit antibodies used were against ALIX (1:1000, SAB4200476**,** Sigma), HA (1:1000, 51064-2-AP, Proteintech), GFP (1:1000, 50430-2-AP), TSG101 (WB) (1:1000, T5701, Sigma), LAMP1 (1:1000, L1418, Sigma). Rabbit anti-*Coxiella* NMII antibodies were generated by Covalab. Rat antibodies against HA were purchased from Roche (1:200, 11867423001). Hoescht 33258, anti-rabbit and anti-mouse HRP-conjugated antibodies were purchased from Sigma. Rat, mouse and rabbit IgG conjugated to Alexa Fluor 488, 555 or 647 as well as Prolong Gold antifade mounting reagent were purchased from Invitrogen. LysoTracker Red (L7528) was purchased from Life Technologies. Purified L-α-phosphatidylcholine (Egg PC) (Cat# 840051P), L-α-phosphatidylserine (Brian PS) (Cat. No. 840032C), lysobisphosphatidic acid (LBPA) (Cat. No. 857133C), N-Rhodamine-PE (Cat. No. 810179) were purchased from Avanti Polar Lipids, Inc. Tetramethylrhodamine (TMR) dextran 70kDa was purchased from Invitrogen (Cat. No. D1818, Invitrogen).

*Transfections, cell cultures and infections*

U2OS, A549, HEK293T, HeLa (ATCC) cells were routinely maintained in DMEM containing 10% (vol/vol) of foetal bovine serum (FBS) in a humidified atmosphere of 5% CO_2_ at 37 °C. For ectopic expression of proteins in mammalian cells, cells were grown to 70% confluence and transfected with JetPEI cationic polymer transfection reagent (Polyplus Transfection) diluted in 150 mM NaCl. SiRNA transfections in cells were performed with calcium phosphate buffer in HEK293T (ATCC) for 24h with siRNA TSG101 (S102664522) and siRNA negative Control (S103650318) from Qiagen. For bacterial infections, U2OS cells were seeded in 24-well plates with coverslip one day prior to infection. Cells were then challenged with different strains at an MOI of 100 followed by 10 minutes of centrifugation at 400 x g for infection synchronization.

*Immunofluorescence staining and microscopy*

Cells were fixed in 4% (wt/vol) paraformaldehyde in phosphate-buffered saline (PBS) pH 7.4 solution at room temperature for 20 min. Samples were then rinsed with PBS solution and incubated in blocking solution (0.5% bovine serum albumin [BSA], 50 mM NH_4_Cl in PBS solution, pH 7.4). Cells were then incubated with the primary antibodies diluted in blocking solution for 1 h at room temperature in a humidified atmosphere, washed five times in PBS solution, and further incubated for 1 h with the secondary antibodies diluted in blocking solution complemented with Hoechst 33258 (Sigma) for DNA staining. Coverslips were then mounted on slides using Fluoromount mounting medium (Sigma)*.* Samples were imaged using an epifluorescence EVOS M5000 (Thermo Fisher Scientific) equipped with 63X or 40X oil immersion objectives. Live cell imaging was performed with a spinning disk Olympus SR microscope equipped with sCMOS Fusion BT Hamamatsu camera. Automated epifluorescence microscopy was performed using an ImageXpress PICO (Molecular Devices) equipped with a CMOS 5Mpx 12bits camera. In this case, 25 fields/well were acquired for image analysis. Image J, ICY and CellProfiler software were used for image analysis and quantifications.

*Legionella pneumophila transformation*

Electrocompetent *L. pneumophila* strain Paris WT or the corresponding Δ*dotB* strain were electroporated with plasmids encoding either BLAM alone or BLAM-Vice using a Gene Pulser™ apparatus (Bio-Rad Laboratories, Richmond, CA) as follows: capacitance, 25 μF, voltage, 2.5 kV, and resistance, 200 Ω. After the pulse was applied, aliquots were immediately resuspended in 2 ml AYE medium and incubated at 37 °C with orbital shaking of 150 rpm for 4 h. The suspension was spread on BCYE plates containing kanamycin (15ug/ml final) and chloramphenicol (5ug/ml final).

*Lentivirus production and stable cell line engineering.*

Lentiviral particles production was performed as previously described (Siadous *et al.* 2020). U2OS cells stably expressing Cas9 were selected using DMEM supplemented with 10% FBS and 20 μg/ml Blasticidin (Invivogen). U2OS cells stably expressing Cas9 and Control- or ALIX-targeting guides were selected using DMEM supplemented with 10% FBS, 20 μg/ml Blasticidin (Invivogen) and 5 μg/ml Puromycin (Invivogen).

*Mutant library screening*

U2OS cells were seeded into black 96-well plates with clear bottom (Greiner Bio-One, 1,200 cells / well) 24h prior to infection. Cells were infected with a sub-library of GFP-tagged *C. burnetii* mutants (MOI of 100) and centrifuged at 400 x g for 10 min at room temperature. Following 45 minutes of incubation at 37°C in a humidified atmosphere of 5% CO_2_, infected cells were washed 3 times in PBS pH 7.4 and DMEM containing 10% FBS was added. After 4 days of infection, cells were washed 3 times with PBS and fixed with PBS pH 7.4 containing 4% PFA. Immunofluorescence using Hoescht 33258 (Sigma), anti-LBPA (Echelon Biosciences) and anti-LAMP1 (Sigma) was performed as described above. 25 fields/well were imaged with an ImageXpress PICO microscope equipped with a CMOS 5Mpx 12bits camera and analyzed with ICY software.

*Beta-Lactamase Translocation Assay*

To assess the translocation of CBU2007 by either *C. burnetii* or *L. pneumophila*, U2OS cells were cultured in black 96-well plates with clear bottom (Greiner Bio-One). The cells were then infected with the appropriate strain at a multiplicity of infection (MOI) of 100 and incubated for 24, 48, and 72 hours with *C. burnetii* or 24 hours with *L. pneumophila* in a humidified atmosphere of 5% CO_2_ at 37 °C. Negative and positive controls were included, consisting of bacteria expressing BLAM alone or T4SS-defective strains (*dotA*::Tn and Δ*dotB*, respectively) expressing either BLAM alone or BLAM-Vice. For *C. burnetii* infections BLAM-tagged CvpB was also used as positive control. For translocation detection, cells were loaded with the fluorescent substrate CCF4/AM from the LiveBLAzer-FRET B/G loading kit (Invitrogen). The loading solution contained 20 mM HEPES, 15 mM probenecid (Sigma) at pH 7.3, diluted in Hank's balanced salt solution (HBSS). After incubating in the dark at room temperature for 1 hour, cells were visualized using an EVOS inverted fluorescence microscope equipped with DAPI and GFP filter cubes. CellProfiler was used to analyze images as previously described[(42)](https://app.readcube.com/library/1811f86c-d2ee-4af8-afcd-e59769f81591/all?uuid=10910450959032347&item_ids=1811f86c-d2ee-4af8-afcd-e59769f81591:f01f3c57-821b-48f0-b6d9-b7f105323583).

*Transmission Electron Microscopy (TEM)*

After 5 days of infection, U2OS cells were immersed in a 2.5% glutaraldehyde solution in PHEM buffer (CliniSciences, pH 7.4) and kept overnight at 4°C. Subsequently, they were rinsed with PHEM buffer and post-fixed with a mixture of 0.5% osmic acid and 0.8% potassium hexacyanoferrate trihydrate in the dark at room temperature for 2 hours. After two additional rinses in PHEM buffer, the cells were dehydrated using a series of ethanol solutions with increasing concentrations (30-100%). The cells were then embedded in EmBed 812 using an Automated Microwave Tissue Processor for Electron Microscopy (Leica EM AMW). Thin sections (70 nm) were obtained from different levels of each block using a Leica-Reichert Ultracut E microtome. These sections were counterstained with 1.5% uranyl acetate in 70% ethanol and 3% lead citrate (Reynolds method - EMS), and observed using a Tecnai F20 transmission electron microscope operating at 120 kV at the Institut des Neurosciences de Montpellier, INSERM U 1298, Université Montpellier, Montpellier, France.

*Correlative Light Electron Microscopy (CLEM)*

U2OS cells seeded on Gridded Coverslips (MATTEK) were transfected 24h prior to image acquisition using an Olympus SR microscope. The following steps of sample preparation were performed at room temperature: cells were fixed in HEPES buffer (pH 7.4) containing 2.5% Glutaraldehyde for 2 hours and rinsed 3 times for 5 minutes with HEPES buffer (pH 7.4). Samples were immersed in the Post Fixation solution (HEPES buffer (pH 7.4) containing 1% osmic acid and 1.5% potassium ferrocyanide) for 1h and rinsed 3 times with HEPES buffer (pH 7.4). The enhancement was performed using a solution of 1% tannic acid in HEPES buffer (pH 7.4) for 30 minutes, then washed 3 times with HEPES buffer (pH 7.4). A second post-fixation treatment was done using 1% osmium acid in distilled water for 1h and washed 3 times with distilled water. For dehydration, samples were immersed in a series of ethanol concentrations: 25%, 50%, 75%, and 95% for 10 minutes each and 3 times 15 minutes in 100% ethanol. The first step of embedding was performed using EPON and 2% BDMA (EMS) overnight, and then areas selected by confocal microscopy were specifically encapsulated with the same solution for at least 2 hours. The resin was polymerized for at least 48 hours at 60°C. Images were acquired with a Tecnai F20 transmission electron microscope operating at 120 kV at the Institut des Neurosciences de Montpellier, INSERM U 1298, Université Montpellier, Montpellier, France.

### *Growth assay using Colony Forming Unit (CFU) assay*

### U2OS cells were plated in duplicate or triplicate in a 6-well plate at 2×10^5^ per well and allowed to adhere overnight. The next day, cells were infected with WT or *vice*::Tn *C. burnetii* in 500 μL DMEM supplemented with 10% FBS for 2 h, then washed 15 times with PBS to remove extracellular bacteria. Cells were scraped into 2 mL fresh DMEM supplemented with 10% FBS and replated in a 24-well plate for analysis at day 3 (500 μL cells), and day 6 (125 μL cells + 325 μL DMEM supplemented with 10% FBS) postinfection. To establish a baseline number of internalized bacteria at day zero, 500 μL of infected cells was centrifuged into a pellet and lysed in sterile water for 5 min. Lysate was serially diluted in ACCM-D and spotted onto ACCM-D agarose plates, with each condition performed in triplicate. ACCM-D agarose plates were incubated for 7 to 10 days at 37°C in 2.5% O_2_ and 5% CO_2_, and the number of colonies was counted to determine bacterial viability.

*Magic red assay for cathepsin B activity*

Active cathepsin B was quantitated in live cells using Magic Red following the manufacturer’s protocol. Briefly, 2x10^5^ U2OS cells were infected with WT or *vice*::Tn *C. burnetii* in 6-well plates for 2 h and washed with PBS. At 1dpi, cells were replated onto ibidi slides (3X10^3^ cells per channel). Magic Red (Cat. 937; ImmunoChemistry Technologies; Bloomington, MN) was reconstituted in 50 μl DMSO, vortexed, and stored at -20°C. Immediately before use, the stock was diluted at 1:10 dilution in sterile water, and then further diluted at 1:25 in DMEM supplemented with 10% FBS. Cells were incubated with 50 μl diluted Magic Red for 30 min at 37°C and 5% CO_2_, washed with pre-warmed media, and Z-stack confocal images obtained with identical capture settings with a Nikon spinning disk confocal microscope at 60X. Images were processed identically with ImageJ, and Magic Red fluorescence intensity normalized to the CCV area. At least 15 cells or CCVs were measured per condition in each of three independent experiments.

*Protein purification*

Codon optimized *vice* was inserted into the pTwist vector and expressed in *Escherichia coli* LEMO21 (New England Biolabs). Bacterial cultures were grown at 37°C to mid-exponential phase (OD600nm  =  0.5) and protein expression was induced overnight at 16°C with 1 mM isopropyl-β-D-thiogalactopyranoside (IPTG, Euromedex). Bacteria were harvested by centrifugation, resuspended in lysis buffer (50 mM Tris pH 7.8, 500 mM NaCl, 15 mM imidazole, complete anti-protease (Roche)) and lysed by sonication with 5 rounds of 10 seconds pulsation at 15 watts and 6 rounds of 10 seconds pulsation at 12 watts with 30 seconds in between each round. Lysates were then cleared by centrifugation (12,000 x g, 30 min, 4°C). Proteins were purified on Econo-Pac Chromatography Columns (Bio-rad) with HIS-Select Nickel Affinity Gel (Sigma). After 5 washes with washing buffer (50 mM Tris pH 7.8, 500 mM NaCl, 50 mM imidazole, complete anti-protease (Roche)) proteins were eluted using elution buffer (50 mM Tris pH 7.8, 500 mM NaCl, 1 M imidazole, complete anti-protease (Roche)).

*Protein–Lipid Overlay Assay*

Purified 6xHIS-GST or 6xHIS-CBU2007 were incubated with PIP Strips membranes (Echelon Biosciences) according to the manufacturer's instructions. Rapidly, TBS-T/BSA (10 mM Tris-HCl pH 8, 150 mM NaCl, 0.1% Tween 20, 3% BSA) was used throughout the assay. Membranes were initially blocked with TBS-T/BSA for 1 hour, then incubated with 1 μg/mL of purified protein overnight at 4 °C. After three washes with TBS-T/BSA, membranes were immunoblotted with anti-HIS HRP-conjugated antibodies for 1 hour at room temperature. Following three washes with TBS-T/BSA, membranes were incubated with ECL for signal detection and visualized using a Chemidoc imaging system (Biorad).

### *In Vitro Cosedimentation Assays with Large Unilamellar Vesicles (LUVs)*

To determine the binding of 6xHIS-CBU2007 to LUVs, different mixtures were made using PC, PS and LBPA at different molar ratios: 100/0/0; 90/10/0; 70/10/20 and 80/0/20. The preparation of lipid mixtures and experiment procedures were performed as previously described[(4)](https://app.readcube.com/library/1811f86c-d2ee-4af8-afcd-e59769f81591/all?uuid=18970346884836142&item_ids=1811f86c-d2ee-4af8-afcd-e59769f81591:a15adb6b-0f02-4ce7-a53d-9f042f1d1bf8).

*Macropinocytosis inhibition assay*

For ectopic expression experiments, U2OS cells were transfected for 8 hours with the appropriate plasmids, washed 3 times with PBS pH 7.4 and incubated overnight with DMEM containing 10% FBS and 50 μM 5-[N-ethyl-N-isopropyl] amiloride (EIPA, MERCK). Cells were then washed 3 times in PBS pH 7.4, fixed in PBS pH 7.4 containing 4% PFA and prepared for fluorescence microscopy as described above. For infection experiments, U2OS seeded on coverslips in 24-well plates were infected with wt *C. burnetii* at an MOI of 100 and incubated 24 h at 37°C and 5% CO_2_. Cells were then incubated with the indicated concentrations of EIPA and further incubated for 48h at 37°C and 5% CO_2_. Cells were then washed in PBS, fixed with PBS containing 4% paraformaldehyde for 20 minutes at room temperature and processed for immunolabeling as previously described.

*Dextran internalization assay*

For ectopic expression experiments, U2OS cells were transfected for 8 hours with the appropriate plasmids, then washed 3 times with PBS pH 7.4 and incubated overnight with DMEM containing 0.5 mg/ml 70 kDa TMR Dextran. Following 3 washes in PBS pH 7.4, cells were imaged with a spinning disk Olympus SR microscope as described above. For infection experiments, U2OS cells were plated in a 6 well plate (2×10^5^ cells per well) and allowed to adhere overnight. Cells were then infected with wt or *vice*::Tn *C. burnetii*, washed extensively with PBS and incubated in 2 ml of fresh growth medium (DMEM 10% FBS). At 1dpi, cells were replated onto ibidi slides (9x10^3^ cells per channel). Dextran was dissolved to a concentration of 25 mg/ml in sterile PBS and stored at -20°C; immediately before use, the stock was diluted to 0.5 mg/ml in DMEM 10% FBS. Cells were incubated with 120 μl diluted dextran overnight at 37°C and 5% CO_2_, washed with pre-warmed media, and Z-stack confocal images were obtained with identical capture settings with a Nikon spinning disk confocal microscope with a 60X objective. Images were processed identically using ImageJ, and dextran fluorescence intensity was normalized to the area of the cell. At least 15 cells were measured per condition in each of three independent experiments.

*N-Rhodamine-PE assay for MVB labeling*

MVBs were labeled with the human fluorescent probe *N*-(lissamine rhodamine B sulfonyl)-phosphatidylethanolamine (N-Rhodamine-PE) according to Savina *et al.*, 2002. Briefly, N-Rhodamine-PE was stored in chloroform/methanol (2:1), dried under nitrogen and solubilized in absolute ethanol to a concentration of 0.4 mM. N-Rhodamine-PE was diluted to 0.5 μM drop by drop into serum-free DMEM using a Hamilton syringe while vortexing. U2OS cells were washed with ice-cold PBS and then incubated in serum-free DMEM containing N-Rhodamine-PE for 1 hour at 4°C. Cells were then extensively washed with ice-cold PBS to remove excess unbound lipids prior to being pulse-chased for 60 minutes at 37°C in serum-free DMEM and fixed with PBS containing 4% paraformaldehyde for 20 minutes at room temperature. Cells were processed for immunolabeling as previously described.

*Virion-Like Particles (VLPs) release assay*

HEK293T cells were transfected with either scramble or TSG101 targeting siRNA for 24h using the RNAiMAX transfection reagent (Thermo Fisher Scientific) according to the manufacturer’s recommendations. Cells were then transfected for 24h with plasmids expressing viral proteins in combination with plasmids expressing either the HA tag alone or HA-Vice. The culture medium of transfected cells was clarified at 800 × g for 5 minutes at 4°C and VLPs were collected after ultracentrifugation at 100,000 × g for 90 minutes at 4°C in an SW41Ti rotor (Beckman Coulter) through a 25% sucrose cushion in TNE buffer (25 mM Tris-HCl, 4 mM EDTA, and 150 mM NaCl). The resulting pellets were resuspended overnight in TNE buffer at 4°C. To assess the release of Gag-VLPs, an anti-Cap24 immunoblot was performed on both supernatant and cell lysate. The calculation for Gag-VLP release was performed using the blot intensity module on Fiji software and applying the following formula: % of Gag in VLP = Gag released/(Gag released + Gag Intracellular normalized to GAPDH).

*Protein co-purification Assay*

LEMO21 bacteria were transformed with pGEX4T1, pGEX4T1-RAB26, pGEX4T1-CHMP3B or pGEX4T1-ALIX (kindly provided by Prof Maryse Lebrun, LPHI UMR5235, Montpellier, France) in combination with pTWIST-Vice. Bacterial cultures were grown at 37°C to mid-exponential phase (OD600nm  =  0.5) and protein expression was induced for 3 hours with 500 μM isopropyl-β-D-thiogalactopyranoside (IPTG, Euromedex). Bacteria were harvested by centrifugation, resuspended in lysis buffer (50 mM Tris pH 7.8, 500 mM NaCl, complete anti-protease (Roche)) and lysed by sonication with 5 rounds of 10 seconds pulsation at 15 watts and 6 rounds of 10 seconds pulsation at 12 watts, with 30 seconds in between each round. Lysates were then cleared by centrifugation (12,000 x g, 30 min, 4°C). Proteins were purified on agarose-glutathione resin (Sigma). After 10 washes with lysis buffer, proteins were eluted using elution buffer (50 mM Tris pH 7.8, 500 mM NaCl, 50 mM reduced L-glutathione (Sigma), complete anti-protease (Roche)).

**Supplementary Figures**

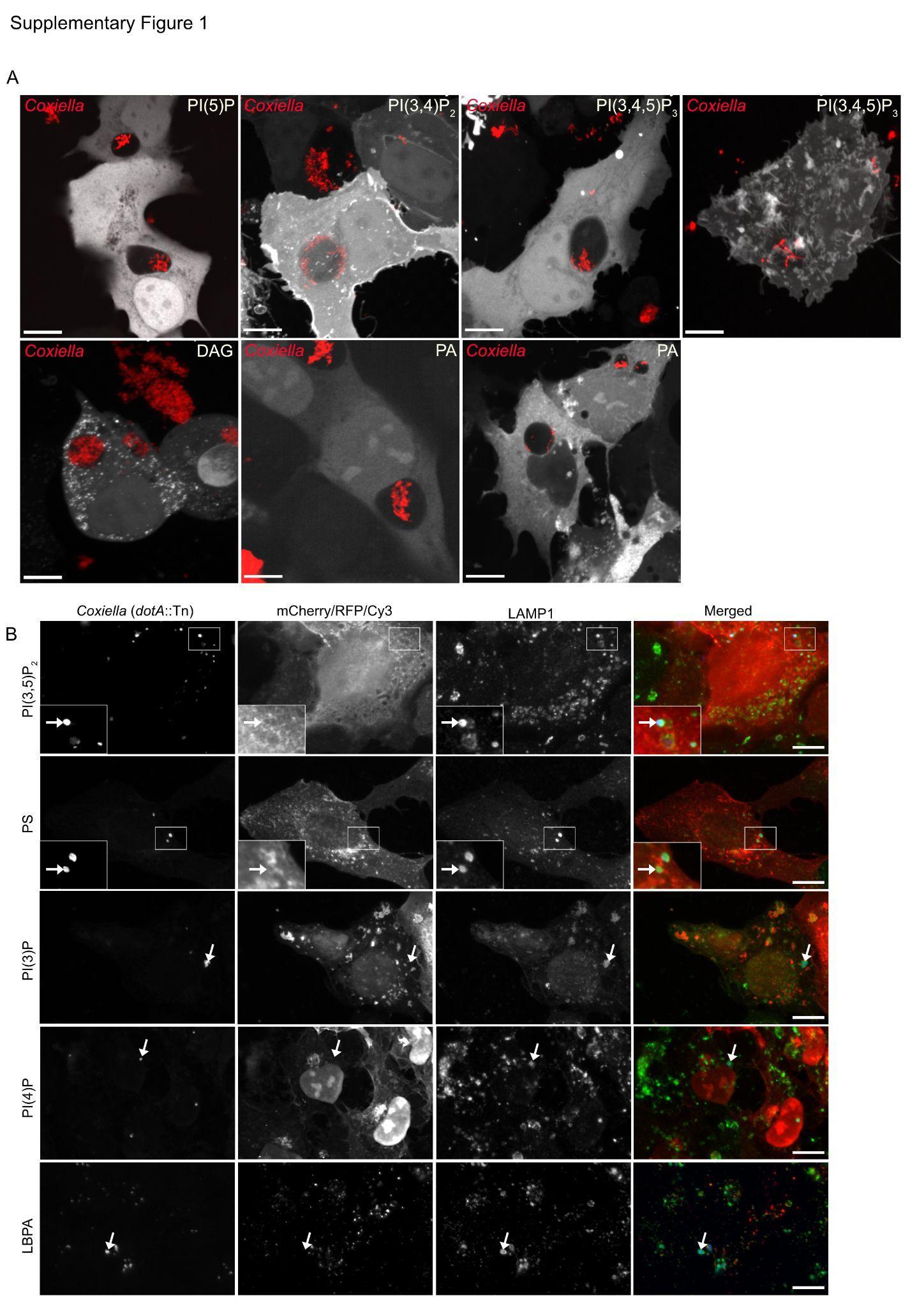

**Supplementary Figure 1: Lipid composition of wt and *dotA::Tn* *Coxiella*-Containing Vacuoles.** (A) U2OS cells were infected with wt *C. burnetii* (red) and transiently transfected with the indicated mCherry/RFP-tagged lipid-binding sensors (white). (B) U2OS cells were challenged for 4 days with the Dot/Icm defective *dotA::Tn* *C. burnetii* mutant strain (blue) and transfected with either mCherry/RFP-tagged lipid-binding sensors or labeled with an anti-LBPA antibody (red). Cells were fixed and stained with an anti-LAMP1 antibody (green). White arrows indicate *C. burnetii* colonies. Scale bars: 10 μm.

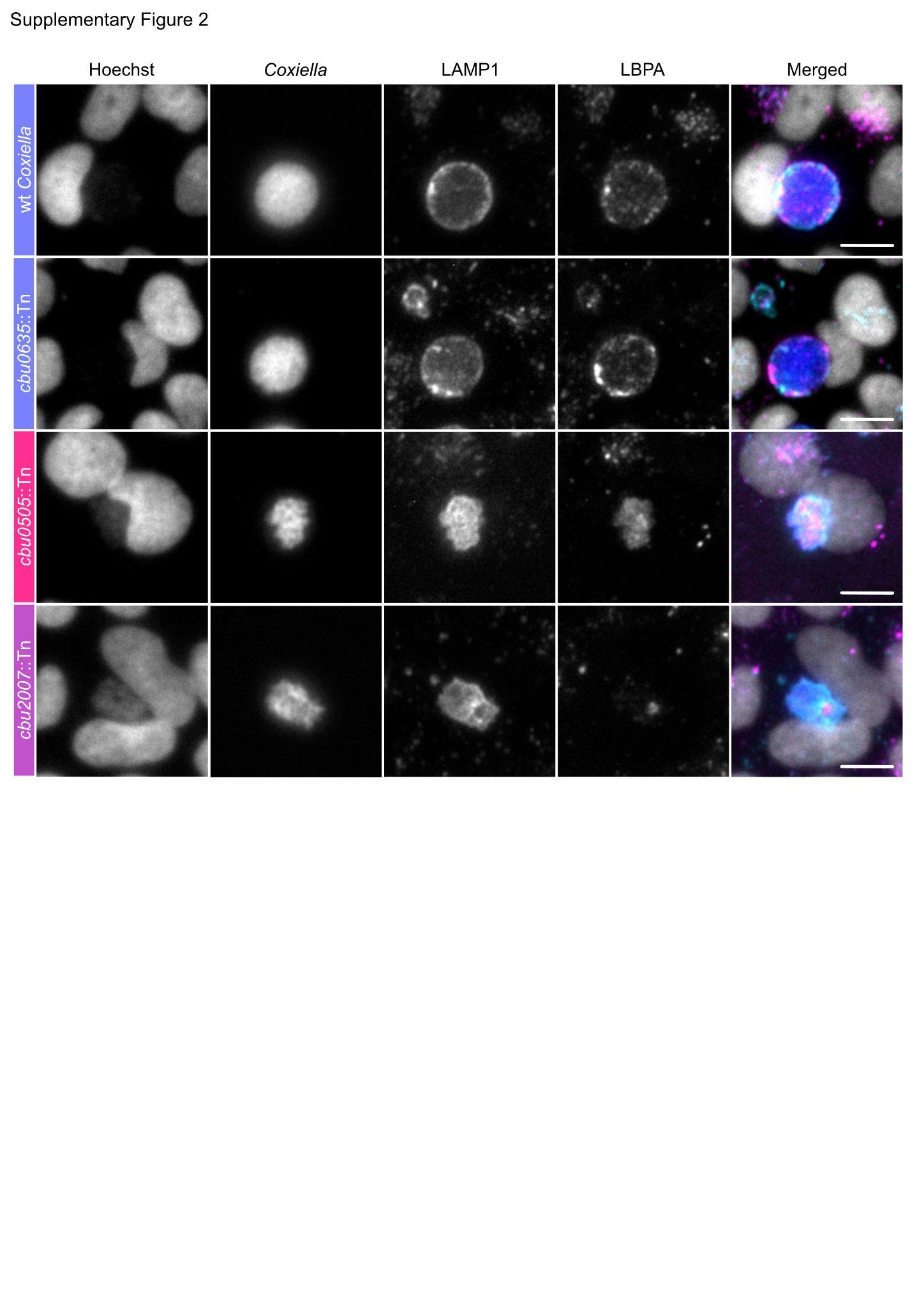

**Supplementary Figure 2: LBPA does not localize to *cbu2007*::Tn mutant strain CCVs.**

Representative images of U2OS cells infected for 4 days either with WT *C. burnetii*, the *cbu0635*::Tn, *cbu0505*::Tn or *cbu2007*::Tn mutant strains. Cells were processed for microscopy using Hoechst (white), anti-LBPA (pink), and anti-LAMP1 (cyan). Scale bars: 10 μm.

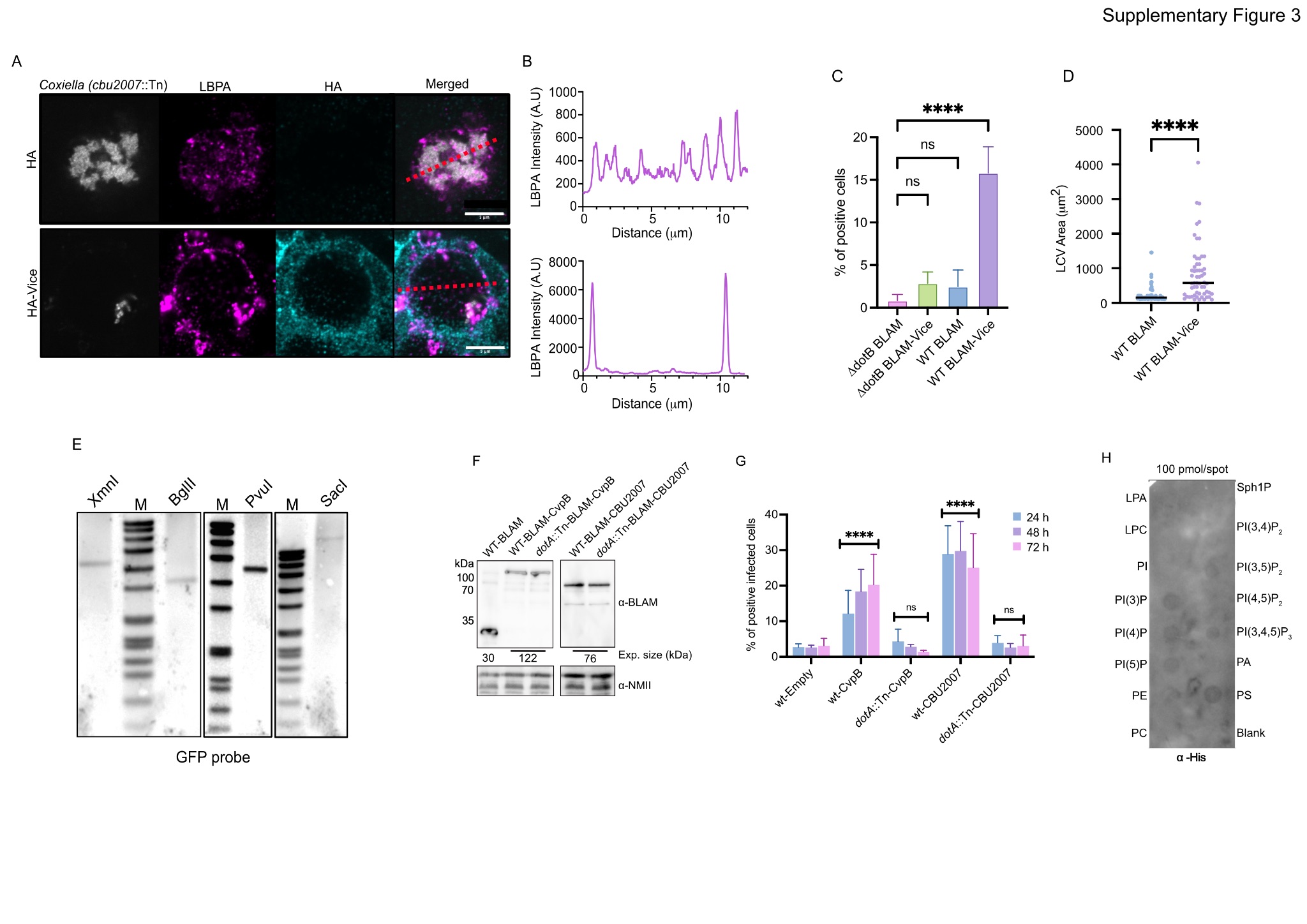
**Supplementary Figure 3: Vice is a T4SS-dependent effector protein required for LBPA recruitment to CCVs.**

(A) U2OS cells were infected for 4 days with the *vice*::Tn *C. burnetii* mutant strain (white) and transfected either with HA alone (upper panel) or HA-Vice for 24 hours (lower panel). Cells were fixed and labeled with anti-LBPA (magenta) and anti-HA (cyan) antibodies. Scale bar: 5 μm. (B) The fluorescence intensity of LBPA (magenta) was measured along the red dash lines illustrated in (A) using ImageJ. (C) U2OS cells were challenged for 24 h either with wt *L. pneumophila* (WT) or the Δ*dotB* mutant strain transformed either with vectors expressing BLAM alone (negative control) or BLAM-Vice. Effector protein translocation was probed using the ꞵ-Lactamase assay. The average percentage of infected cells positive for cleaved CCF-4 was automatically calculated using CellProfiler over the total number of infected cells per each condition. Values are mean ± SD from 3 independent experiments (n.s.: non-significant, ****P<0.0001, one-way ANOVA, Bonferroni’s multiple comparison test). (D) The area of *L. pneumophila*-containing vacuoles (LCVs) was measured by CCF-4 exclusion from U2OS cells infected with WT *L. pneumophila* expressing either BLAM alone or BLAM-Vice. Values are mean ± SD from 3 independent experiments where 50 LCVs were measured for each condition (****P<0.0001, Unpaired t-test). (E) Southern blot analysis of the *cbu2007*::Tn *C. burnetii* mutant strain *Tn1577* genomic DNA digested either with XmnI, BglII, PvuI or SacI and probed with a GFP probe. The unique band observed confirms the unique insertion of the transposon in the *cbu2007*::Tn mutant genome. (F) Lysates of wt *C. burnetii* GFP (WT) or the *dotA*::Tn mutant strain transformed with pXDC61K vectors expressing BLAM alone, BLAM-CvpB or BLAM-CBU2007/Vice were probed using anti-BLAM and anti-*C. burnetii* (NMII) antibodies by Western blot. The expected size of BLAM-tagged proteins is indicated below the top blot. (G) U2OS cells were challenged for 24 h, 48 h or 72 h either with wt *C. burnetii* GFP (WT) or the *dotA*::tn *C. burnetii* mutant strain transformed either with vectors expressing BLAM alone (negative control), BLAM-CvpB (positive control) or BLAM-CBU2007/Vice. Effector protein translocation was assessed as described in C. Values are mean ± SD from 3 independent experiments (n.s.: non-significant, ****P<0.0001, one-way ANOVA, Bonferroni’s multiple comparison test). (H) Protein-lipid overlay assay performed with 6xHIS-GST as the control for the overlay illustrated in Fig 2A.

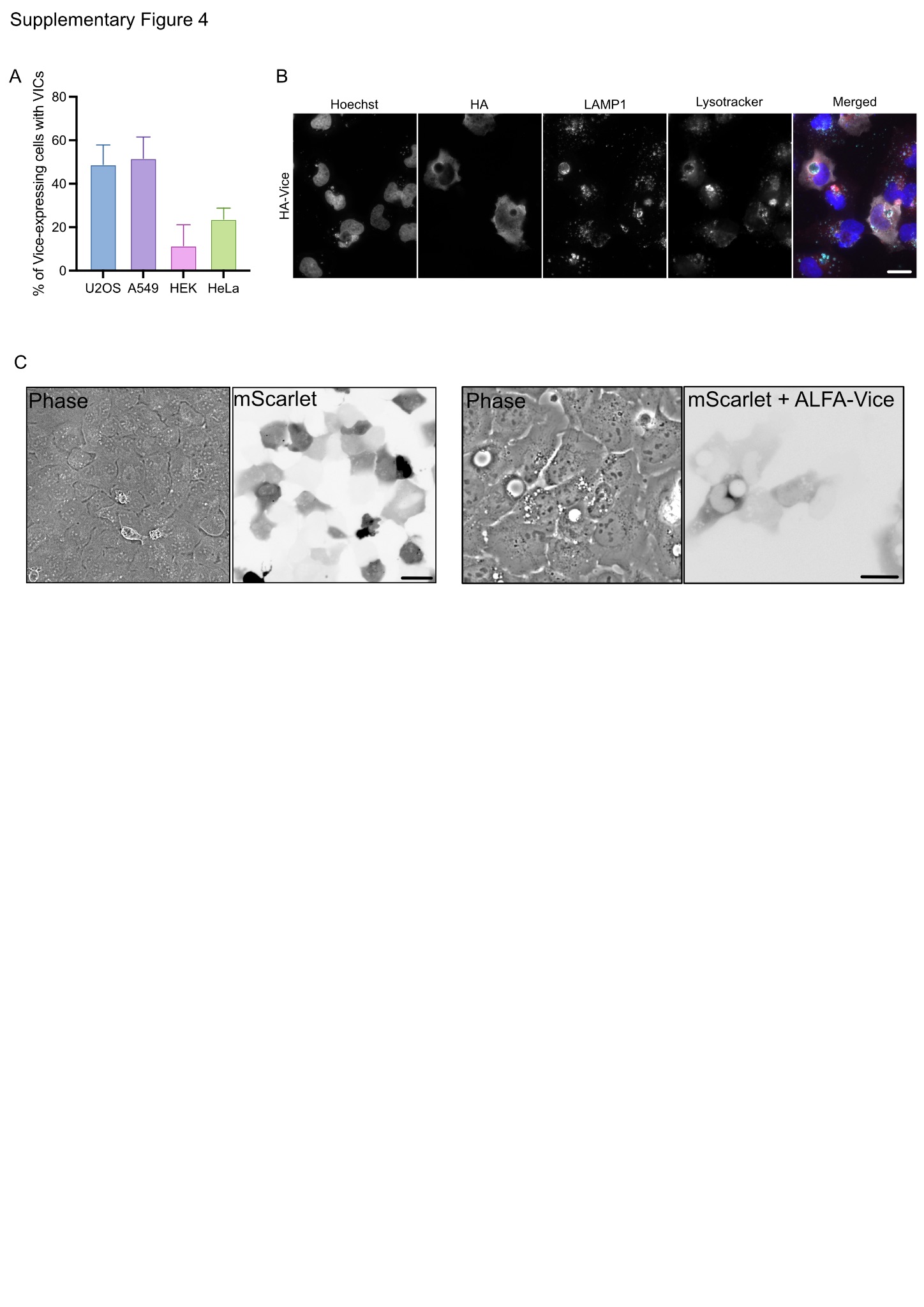

**Supplementary Figure 4: Occurrence and characterisation of Vice-induced vacuoles (VICs).**

(A) Occurrence of VICs was assessed from U2OS, A549, HEK and HeLa cells transfected with plasmids encoding HA-Vice, fixed and labeled with anti-LAMP1 antibodies. Values are mean ± SD from 3 independent experiments where 200 cells were scored for each condition. (B) U2OS cells transfected with a plasmid encoding HA-Vice were incubated with 1 μM Lysotracker (red) for 1 hour, fixed and immunolabeled with anti-HA (white) and anti-LAMP1 (cyan) antibodies. Nuclei were labeled using Hoechst (blue). Scale bars, 20 μm. (C) U2OS cells transfected with a plasmid encoding NbALFA-mScarlet alone or in combination with HA-ALFA-Vice for 8 hours prior to images acquisition in phase contrast (left panel) and 555 nm emission (right panel) channels. Scale bars, 20 μm.

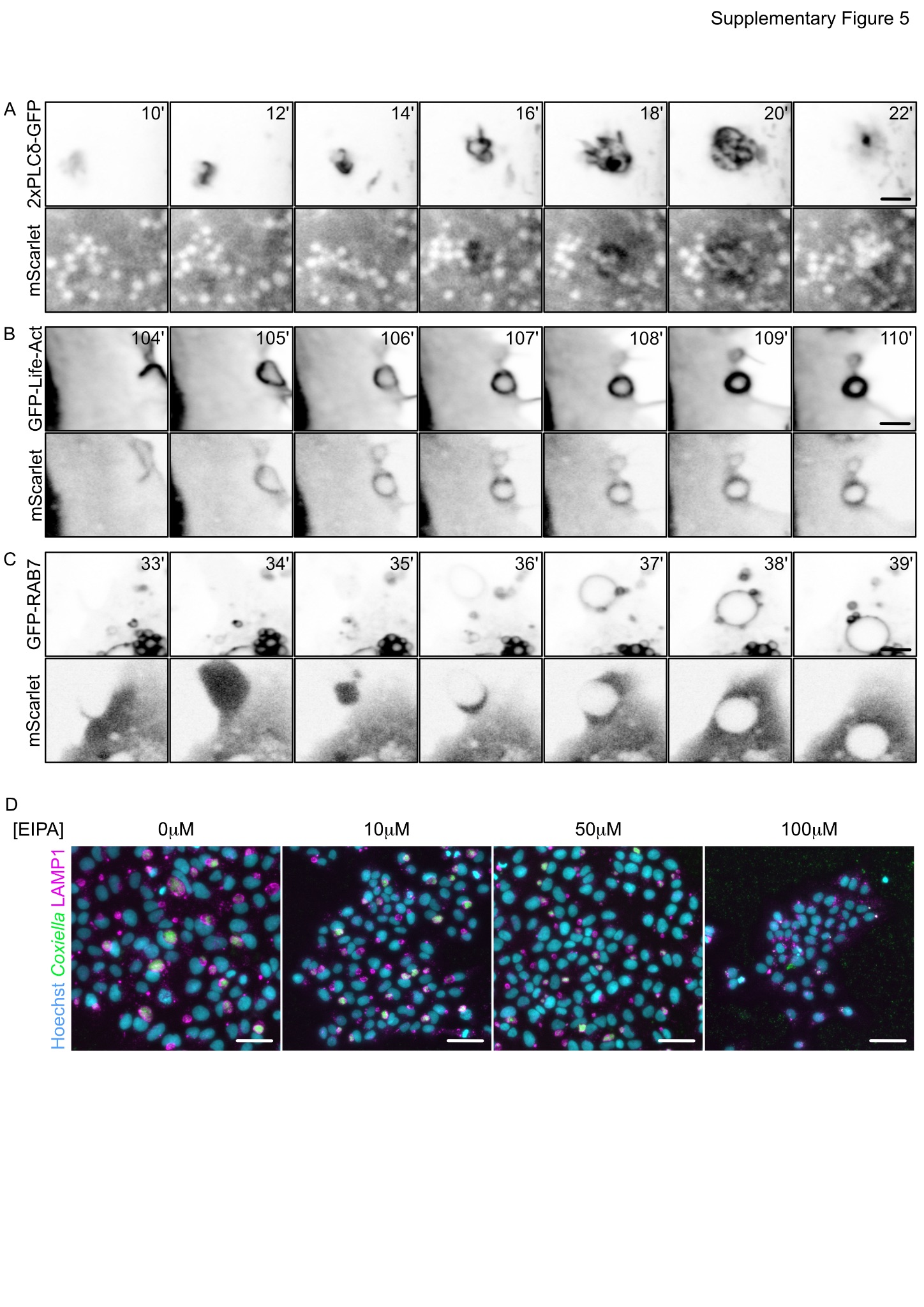

**Supplementary Figure 5**: **Vice stimulates macropinocytosis.**

(A-C) Representative images from Movies S3 to S5 illustrating U2OS cells co-transfected with plasmids encoding HA-ALFA-Vice, NbALFA-mScarlet and either 2xPLCδ-GFP (A, Movie S3), Lifeact-GFP (B, Movie S4) and GFP-RAB7 (C, Movie S5). After 8 hours post-transfection, images were acquired every minute overnight in 555 nm emission (lower panel) and 488 nm emission (upper panel) channels. Scale bars: 4 µm. (D) Representative images of U2OS cells infected with WT *C. burnetii* (green) and incubated with the indicated concentrations of EIPA. 3 days post infection cells were fixed and labeled with anti-LAMP1 antibodies (magenta) and Hoechst (cyan). Scale bars 40 µm.

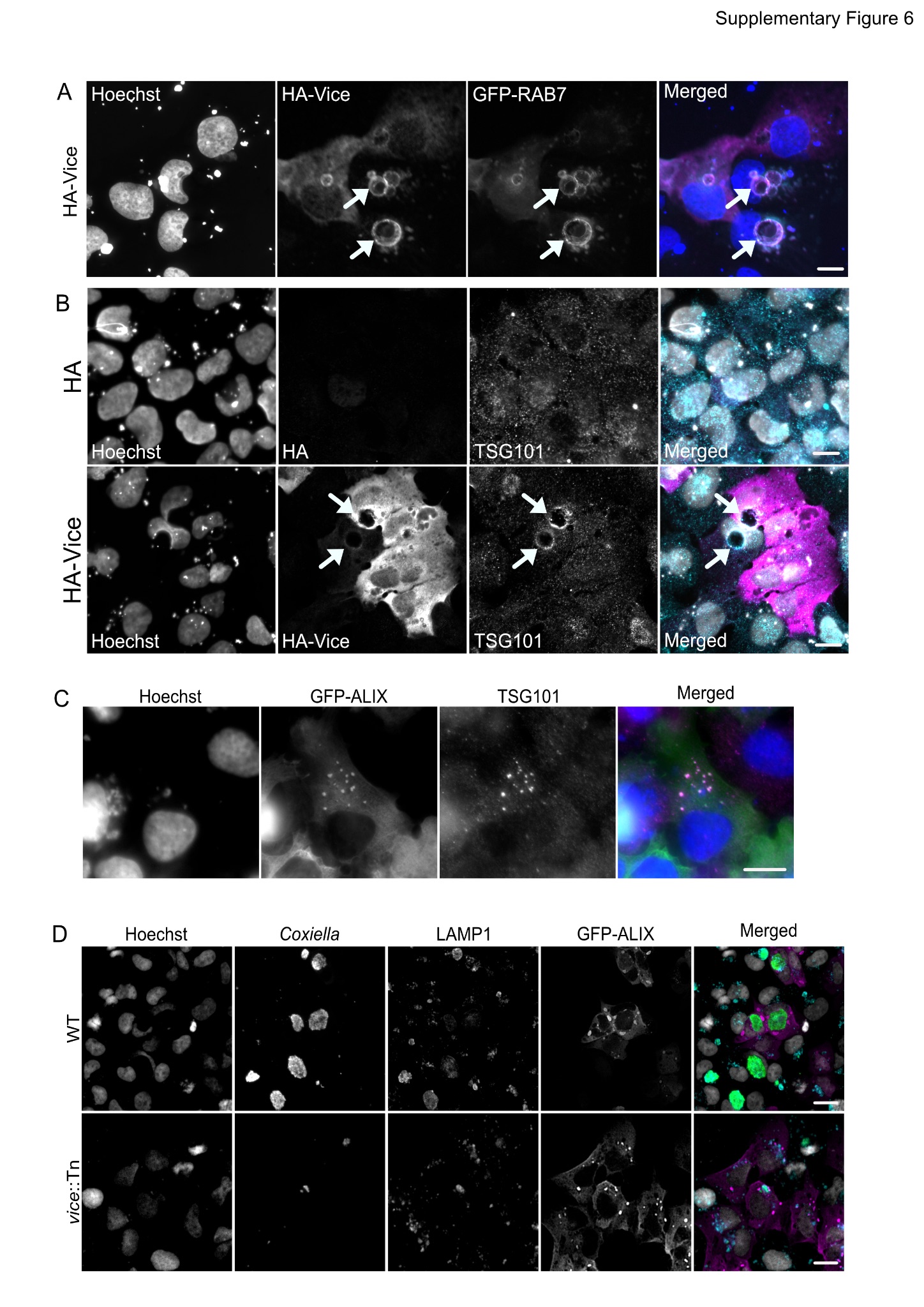

**Supplementary Figure 6**: **Vice subversion of the ESCRT machinery**

(A) U2OS cells transfected with plasmids encoding HA-Vice (magenta) and GFP-RAB7 (cyan) indicate that RAB7 localizes to VICS (white arrows). (B) U2OS cells were transfected with a plasmid encoding HA (upper panels) or HA-Vice (lower panels) and labeled with anti HA (magenta) and anti-TSG101 antibodies (cyan). White arrows point at VICs, scale bars: 10 µm. (C) U2OS cells transfected with a plasmid encoding GFP-ALIX were fixed 24 hours post transfection and labeled with anti-TSG101 antibodies. Scale bar 10 µm. (D) Representative images of U2OS cells infected for 4 days with either WT *C. burnetii* or the *vice*::Tn mutant and transfected for 24h with a plasmid encoding GFP-ALIX (magenta). After fixation, cells were labeled with Hoechst (white), anti-*C. burnetii* (green) and anti-LAMP1 antibodies (cyan). Scale bars: 20 µm. In all conditions, Hoechst was used to label nuclei (white, blue).

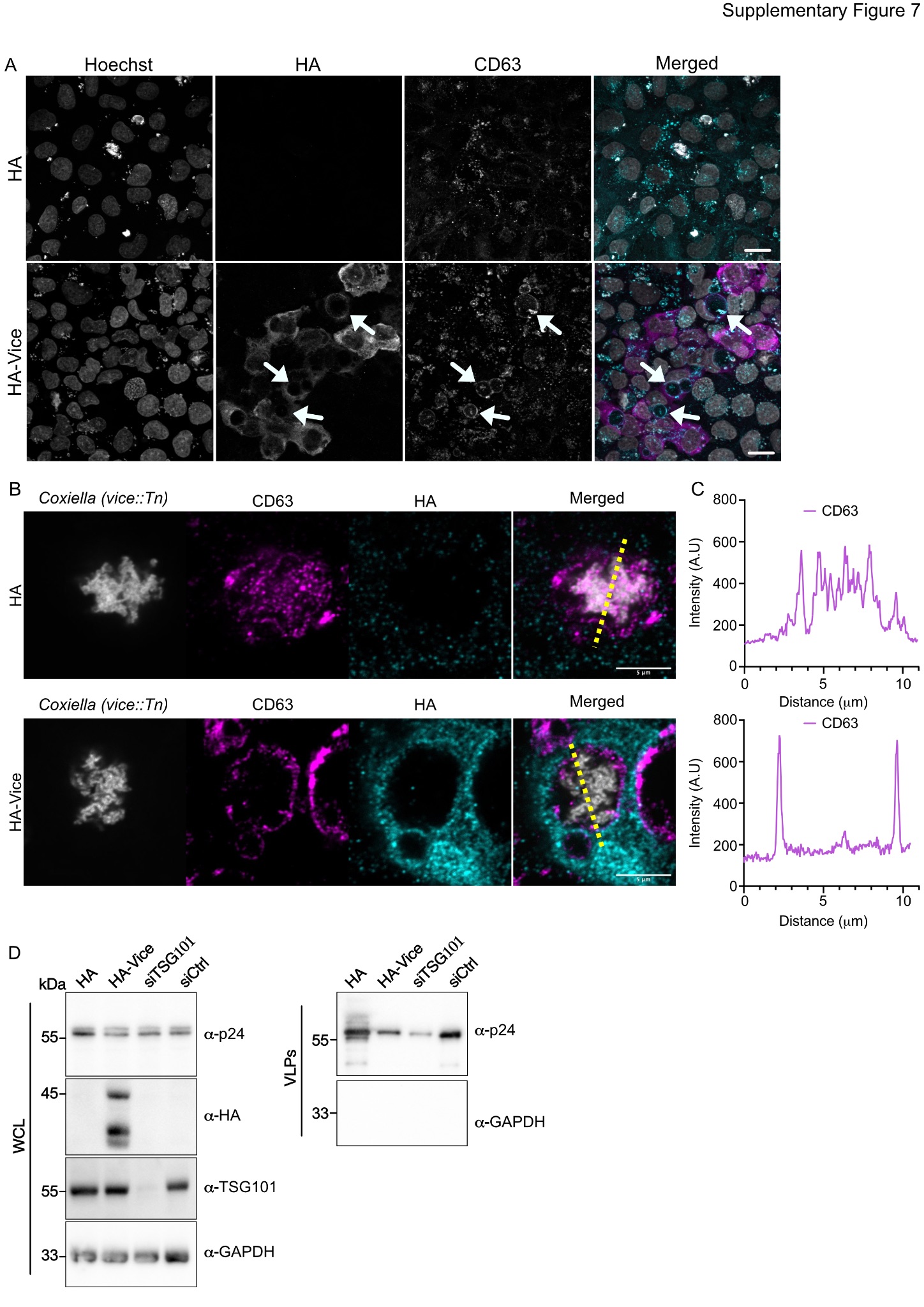

**Supplementary Figure 7: Vice subversion of the ESCRT machinery**

(A) U2OS cells were transfected with plasmids encoding either HA or HA-Vice (magenta) and labeled with an anti-CD3 antibody (cyan). CD63 localizes to VICs (white arrows). Hoechst was used to label nuclei (white). Scale bar 10 µm. (B) U2OS cells were infected with the *vice*::Tn mutant strain (white) for 4 days and transfected with plasmids encoding either the HA tag alone (upper panels) or HA-Vice (lower panels) for 24 hours. Cells were fixed and labeled with anti-CD63 (magenta) and anti-HA (cyan) antibodies. Scale bar 5 µm. (C) Representative distribution of CD63 (magenta) fluorescence intensity was measured along the corresponding yellow dashed lines illustrated in (B). (D) HEK293T were co-transfected with plasmids encoding either HA or HA-Vice and GAG, or transfected with plasmids encoding GAG, 24 hours after transfection with either TSG101-targeting siRNAs (siTSG101) or scrambled siRNA (siCTRL). Virus-like particles (VLP) release was assessed by Western blot using anti-p24 antibodies and the relative percentage of VLPs was measured using ImageJ. Anti-HA, anti-TSG101 and anti-GAPDH were used as controls (WCL: whole cell lysate).

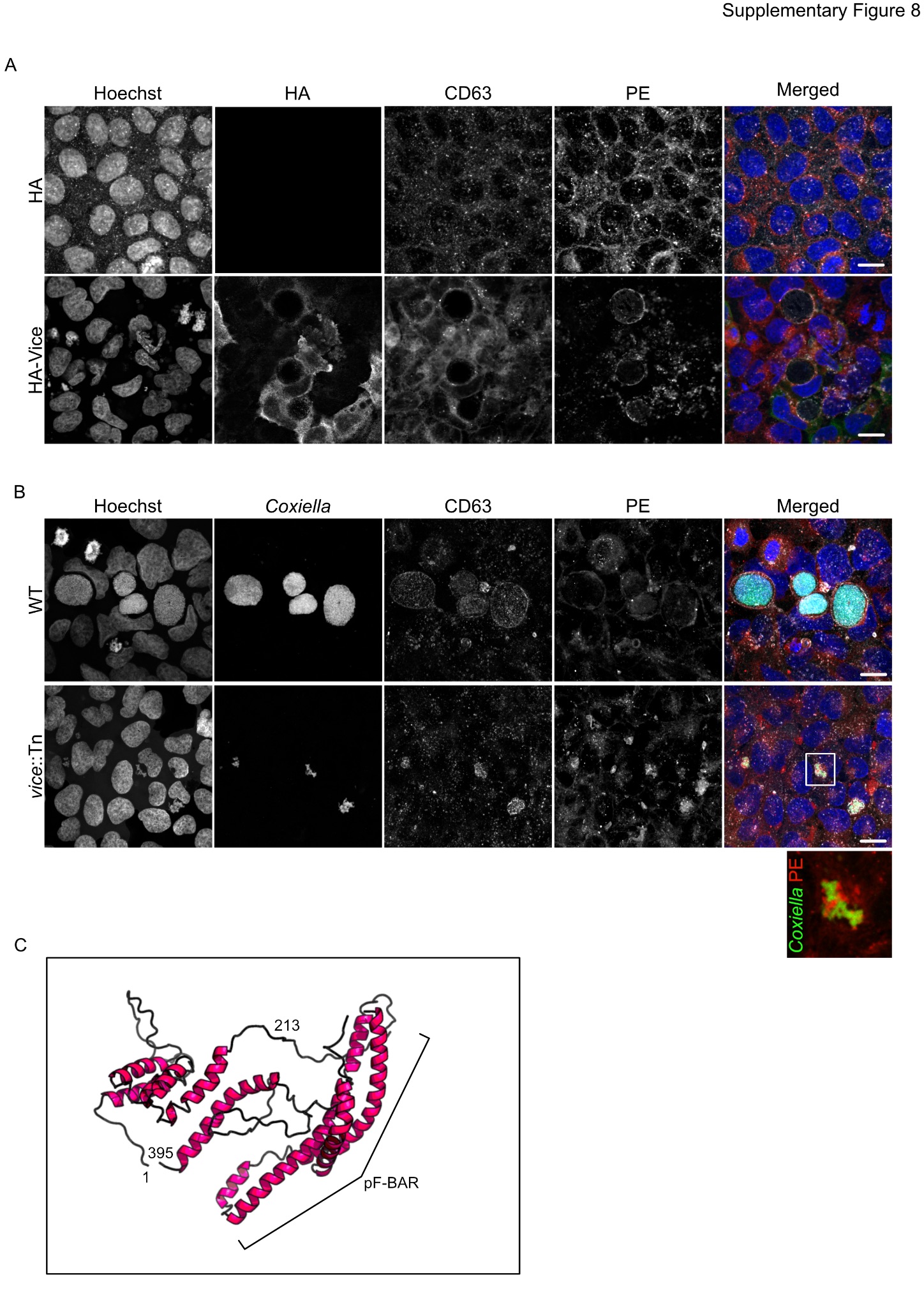

**Supplementary Figure 8: Vice perturbs ILV biogenesis**

(A) U2OS cells transfected with plasmid encoding the HA tag alone (upper panels) or HA-Vice (lower panels, green) were loaded with Rhodamine-PE (red) for one hour, chased for another hour, fixed and labeled with anti-CD63 antibodies (white) and Hoechst (blue). Scale bar 10 µm. (B) U2OS cells were infected with WT *C. burnetii* or the *vice*::Tn mutant strain (green) for 4 days, loaded with Rhodamine-PE (red) for one hour, chased for another hour, fixed and labeled with anti-CD63 antibodies (white) and Hoechst (blue). Scale bars 10 µm. (C) Secondary structure prediction of Vice illustrating the predicted F-BAR domain (pF-BAR).

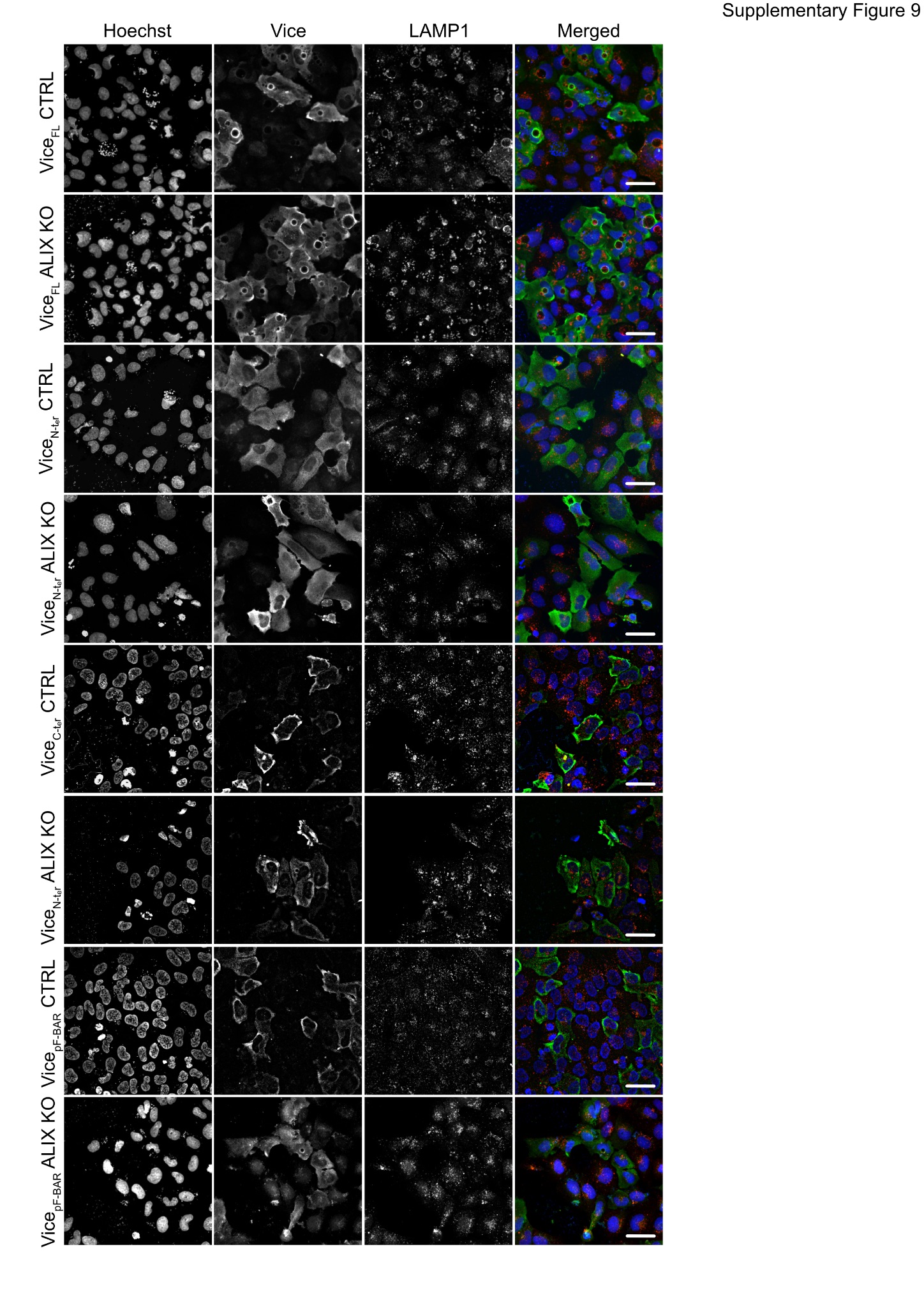

**Supplementary Figure 9: Functional prediction of Vice domains**

(A) U2OS Cas9 Control (CTRL) or ALIX KO cells expressing the indicated HA-tagged Vice truncations were fixed 24 hours post transfection and labeled with anti HA (green), anti-LAMP1 (red) antibodies and Hoechst (blue). Scale bars 20 µm.

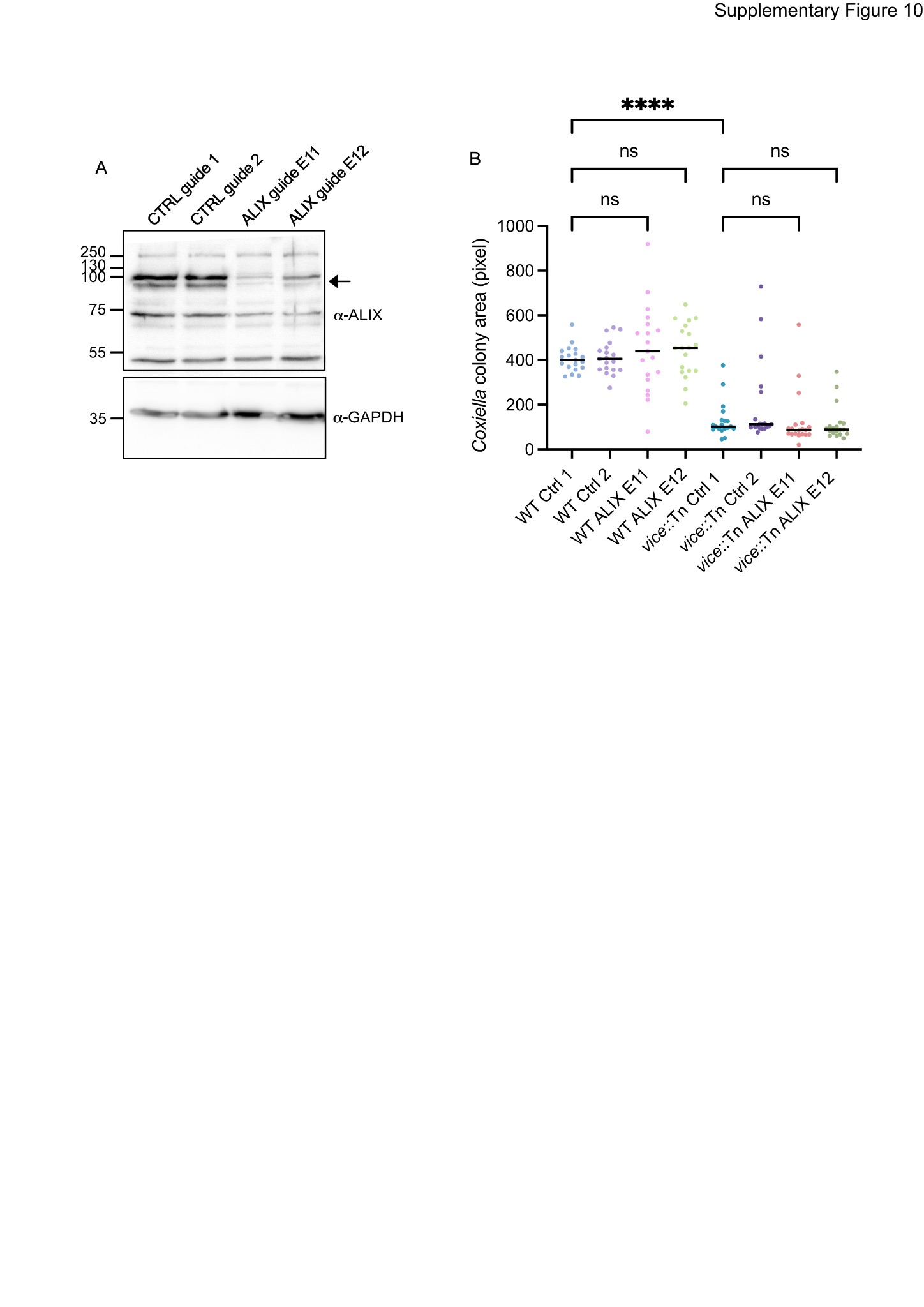

**Supplementary Figure 10: Generation and test of ALIX-KO cell lines**

Immunoblot against ALIX of lysates from U2OS Cas9 cells transduced with lentiviral vectors expressing non-targeting guides (CTRL guide 1 and CTRL guide 2) or ALIX-targeting guides (ALIX guide E11 and ALIX guide E12). Anti-GAPDH was used as loading control. (B) The median area of *C. burnetii* colony area was automatically measured from images of U2OS cells from A, infected with either WT *C. burnetii* or the *vice*::Tn mutant strain for 4 days, fixed and labeled with anti-LAMP1 antibodies. Values are means from 3 independent experiments (n.s.: non-significant, ****P<0.0001, one-way ANOVA, Bonferroni’s multiple comparison test).

**Supplementary movies legends**

**Movie S1**: U2OS cells co-transfected with plasmids encoding HA-ALFA-Vice and NbALFA-mScarlet were imaged after 8 hours in phase contrast (left panel) and 555 nm emission (right panel) channels. Images were acquired every minute for 2h and 20 min.

**Movie S2**: U2OS cells co-transfected with plasmids encoding HA-ALFA-Vice and NbALFA-mScarlet were imaged after 24 hours in phase contrast channels. Images were acquired every minute for 10h and 33 min.

**Movie S3**: U2OS cells co-transfected with plasmids encoding the PI(4,5)P_2_ sensor 2xPLCδ-GFP (left panel, cyan), HA-ALFA-Vice and NbALFA-mScarlet (middle panel, magenta) were imaged after 8 hours. Images were acquired every minute for 26 min.

**Movie S4**: U2OS cells co-transfected with plasmids encoding Lifeact-GFP (left panel, cyan), HA-ALFA-Vice and NbALFA-mScarlet (middle panel, magenta) were imaged after 8 hours. Images were acquired every minute for 4h and 43 min.

**Movie S5**: U2OS cells co-transfected with plasmids encoding GFP-RAB7 (left panel, cyan), HA-ALFA-Vice and NbALFA-mScarlet (middle panel, magenta) were imaged after 8 hours. Images were acquired every minute for 1h and 30 min.

**Movie S6**: U2OS cells co-transfected with plasmids encoding HA-ALFA-Vice and NbALFA-mNeonGreen (middle panel, magenta) for 8 hours were incubated overnight with TMR-Dextran (cyan, left panel). Images were acquired every minute for 2h and 28 min.

**Movie S7**: U2OS cells co-transfected with plasmids encoding the PI(4,5)P_2_ sensor 2xPLCδ-GFP (left panel, red), HA-ALFA-Vice_C-ter_ and NbALFA-mScarlet (middle panel, green) were imaged after 8 hours. Images were acquired every minute for 140 min.

**Movie S8**: U2OS cells co-transfected with plasmids encoding the PI(4,5)P_2_ sensor 2xPLCδ-GFP (left panel, red), HA-ALFA-Vice_p-F-BAR_ and NbALFA-mScarlet (middle panel, green) were imaged after 8 hours. Images were acquired every minute for 151 min.

**Movie S9**: U2OS cells co-transfected with plasmids encoding the PI(4,5)P_2_ sensor 2xPLCδ-GFP (left panel, red), HA-ALFA-Vice_N-ter_ and NbALFA-mScarlet (middle panel, green) were imaged after 8 hours. Images were acquired every minute for 201 min.

**Supplementary Tables**

**Table S1. List of lipid-binding sensors and markers used in this study**

| **Sensor/marker** | **Lipid** | **Source** |
| --- | --- | --- |
| **2xFYVE** | **PI(3)P** | pmCherry-2xFYVE was a gift from Harald Stenmark (Addgene plasmid # 140050 ; http://n2t.net/addgene:140050 ; RRID:Addgene_140050) |
| **OSBP-PH** | **PI(4)P** | pBGPa-CMV-GFP-OSBP PH was a gift from Tim Levine (Addgene plasmid # 58724 ; http://n2t.net/addgene:58724 ; RRID:Addgene_58724) |
| **2xPHD** | **PI(5)P** | GFP-2xPHD was a gift from Bernard Payrastre (Université de Toulouse) |
| **Akt-PH** | **PI(3,4)P_2_/**  **PI(3,4,5)P_3_** | GFP-C1-AKT-PH was a gift from Tobias Meyer (Addgene plasmid # 21218 ; http://n2t.net/addgene:21218 ; RRID:Addgene_21218) |
| **ML1N-PH** | **PI(3,5)P_2_** | ML1Nwt_in_pEGFP-C1 was a gift from Rob Parton (Addgene plasmid # 67797 ; http://n2t.net/addgene:67797 ; RRID:Addgene_67797) |
| **2xPLCδ-PH** | **PI(4,5)P_2_** | 2PH-PLCdelta-GFP was a gift from Sergio Grinstein (Addgene plasmid # 35142 ; http://n2t.net/addgene:35142 ; RRID:Addgene_35142) |
| **BTK-PH** | **PI(3,4,5)P_3_** | PH-Btk-GFP was a gift from Tamas Balla (Addgene plasmid # 51463 ; http://n2t.net/addgene:51463 ; RRID:Addgene_51463) |
| **Gab2-PH** | **PI(3,4,5)P_3_** | PH-Gab2-GFP was a gift from Sergio Grinstein (Addgene plasmid # 35147 ; http://n2t.net/addgene:35147 ; RRID:Addgene_35147) |
| **Lact-C2** | **PS** | mRFP-Lact-C2 was a gift from Sergio Grinstein (Addgene plasmid # 74061 ; http://n2t.net/addgene:74061 ; RRID:Addgene_74061) |
| **PKCɣ-C1A** | **DAG** | GFP-C1-PKCgamma-C1A was a gift from Tobias Meyer (Addgene plasmid # 21205 ; http://n2t.net/addgene:21205 ; RRID:Addgene_21205) |
| **Spo1op/Spo2op** | **PA** | GFP-Spo1op and GFP-Spo2op were gifts from Nicolas Vitale (University of Strasbourg, FR) |
| **anti-LBPA** | **LBPA** | Purified Mouse Anti-LBPA antibodies were purchased from Echelon Biosciences (cat. N. Z-PLBPA) |

**Table S2. List of strains used in this study**

| **Strain** | **Description** | **Origin** |
| --- | --- | --- |
| **WT *C. burnetii*** | *C. burnetii* RSA 439 Nine Mile II carrying a transposon insertion between *cbu1847b* and *cbu1849* (*Tn1832*) | Our Laboratory (Martinez *et al.* 2014) |
| ***C. burnetii Tn1577*** | *C. burnetii* RSA 439 Nine Mile II carrying a transposon insertion GFP-CAT in *cbu2007* | Our laboratory |
| ***C. burnetii Tn292*** | *C. burnetii* RSA 439 Nine Mile II carrying a transposon insertion GFP-CAT in *cbu1648* (*dotA*) | Our Laboratory (Martinez *et al.* 2014) |
| **LEMO21(DE3)** | *E. coli strain used for T7-driven recombinant protein expression* | New England Biolabs |
| ***Legionella pneumophila* Paris strain** | *Legionella pneumophila* Paris wild type strain | Gift from N. Personnic, CIRI Lyon, France |
| ***Legionella pneumophila* Paris strain Δ*dotB*** | *Legionella pneumophila* Paris strain deleted for the gene *dotB* | Gift from N. Personnic, CIRI Lyon, France |

**Table S3. List of oligonucleotides used in this study**

| **Primer Name** | **Sequence** |
| --- | --- |
| **BLAM Translocation Assay** | |
| **Generation of pXDC61K-BLAM derivatives** | |
| CBU2007-BamHI | AGGGGATCCATGACAGTTTATTCCCGATATGA |
| CBU2007-XbaI-rev | AGGTCTAGACTAAAGGGTCCGGTATTTGAAG |
| **Ectopic Expression** | |
| **Generation of pLVX-mCherry-N2 derivatives** | |
| PH-PLCδ-MluI | aggacgcgtatggactcgggccgg |
| PH-PLCδ-NotI-rev | agggcggccgctcgcatccatggagcctgagtggtg |
| PH-Btk-MluI | aggacgcgtatgcagaaagaagaagctatggcc |
| PH-Btk-NotI-rev | agggcggccgctcgaggttttaagcttccattcctgtt |
| **Generation of pLVX-mCherry-C1 derivatives** | |
| AKT-PH-EcoRI | agggaattcaaacgacgtagccattgtgaag |
| AKT-PH-BamHI-rev | aggggatccctatgaattccatggtcacacgg |
| PH-Gab2-EcoRI | agggaattcaatgagcggcggcggc |
| PH-Gab2-BamHI-rev | aggggatccttagaagccgcagatctggc |
| **Generation of mCherry2-C1 derivatives** | |
| OSBP-PH-HindIII | AAGCTTcatcggctcgagaggg |
| OSBP-PH-XmaI-rev | CCCGGGtcacgaattcttcttcacagct |
| ML1N-XhoI | CTCGAGGGtactctgacctgactatggcaacac |
| ML1N-XmaI-rev | aggtctagattaattatccaggtcagggggg |
| PKCg-XhoI | CTCGAGctcacaagttcaccgctcg |
| PKCgamma-HpaI-rev | GTTAACctagatccggtggatccc |
| **Generation of pRK5-HA derivatives** | |
| CBU2007-BamHI-Fw | CCAGGATCCACAGTTTATTCCCGATATGAATTTC |
| CBU2007-HindIII-rev | GCTAAGCTTCTAAAGGGTCCGGTATTTGAAG |
| CBU2007_213_-HindIII-Rv | GCTAAGCTTCTAAGTTTCTTCAGGAACAAAATAGGTG |
| CBU2007_214_-BamHI-Fw | CCAGGATCCATGAAAGTTCAGAATTATGTTCGGTC |
| CBU2007_345_-HindIII-Rv | GCTAAGCTTCTATTCTGGGTTAGTTTTTATTGAATCAATTTTAG |

| **Generation of pRK5-HA-ALFA** | |
| --- | --- |
| MluI-ALFA-Fw | CAAACGCGTATGCCATCACGTTTGGAAGAGGAACTGAGACGCCGCTTAACTGAACCTGGATCCCGGGTCGCGAATTC |
| MluI-pRK5-HA-Rv | CAAACGCGTCGCGCCAGCGTAATCTGGAAC |

| **Generation of pCMV-NbALFA-mScarlet and pCMV-NbALFA-mNeonGreen** | |
| --- | --- |
| Nanobody-EcoRI-Fw | GTAGAATTCTGGCTCTGGTGATGCATCTG |
| Nanobody-BamHI-Rv | GTTGGATCCTCAAGAAGTTTGTGGTTTTGGTGTCTTG |

| **Generation of pET28a-GST** | |
| --- | --- |
| GST-NheI-Fw | GAGTCAGCTAGCATG TCC CCT ATA CTA GGT TAT TGG |
| GST-BamHI-Rv | CAAGGATCCAACCAGATCCGATTTTGGAG |

| **Generation of pGEX4T1 derivatives** | |
| --- | --- |
| RAB26-BamHI-Fw | GAAGGATCCatgtccaggaagaagaccc |
| RAB26-EcoRI-Rv | GTCGAATTCTCAAGGGCGGCAGCAGGAG |
| CHMP3-XmaI-Fw | GCTCCCGGGTGCTGGGCTGTTTGGAAAGACCC |
| CHMP3-NotI-Rv | AGCGCGGCCGCAGCCTAGCTGCGGAGTGTGGC |

**Table S4. List of plasmids used in this study**

| **Name** | **Description** | **Origin** |
| --- | --- | --- |
| **pXDC61K** | IPTG-inducible expression vector for N-terminal fusion of Beta-lactamase (BLAM) in *C. burnetii* | (Burette *et al.* 2020) |
| **pXDC61K-CBU0021** | IPTG-inducible expression of BLAM-CBU0021 (CvpB) | (Martinez *et al.* 2016) |
| **pXDC61K-CBU2007** | IPTG-inducible expression of BLAM-CBU2007 | This study |
| **pLVX-mCherry-N2** | CMV expression vector expressing a C-terminal mCherry tag | (Martinez *et al.* 2016) |
| **mCherry2-C1** | CMV expression vector expressing a N-terminal mCherry tag | Addgene plasmid # 54563 Michael Davidson |
| **pCL-Neo-mCherry-2xFYVE** | Encodes mCherry-2xFYVE (PI3P probe) | (Martinez *et al.* 2016) |
| **mCh2-C1-OSBP-PH** | Encodes mCherry-OSBP-PH (PI4P probe) | This study |
| **mCherry-PHD2X** | Encodes mCherry-2xPHD (PI5P probe) | Bernard Payrastre |
| **pLVX-mCherry-AKT-PH** | Encodes mCherry-AKT-PH (PI3,4P_2_ probe) | This study |
| **mCh2-C1-ML1N-PH** | Encodes mCherry-ML1N-PH (PI3,5P_2_ probe) | This study |
| **pLVX-PLCδ-2xPH-mCherry** | Encodes PLCδ-2xPH-mCherry (PI4,5P_2_ probe) | This study |
| **pLVX-mCherry-Gab2-PH** | Encodes mCherry-Gab2-PH (PI3,4,5P_3_ probe) | This study |
| **pLVX-BTK-PH-mCherry** | Encodes mCherry-BTK-PH (PI3,4,5P_3_ probe) | This study |
| **mRFP-Lactadherin-C2** | Encodes RFP-Lactadherin-C2 (PS probe) | Addgene # 74061 Sergio Grinstein |
| **GFP-Lactadherin-C2** | Encodes GFP-Lactadherin-C2 (PS probe) | Addgene #22852 Sergio Grinstein |
| **mCh2-C1-PKCɣ-PH** | Encodes mCherry-PKCɣ-PH (DAG probe) | This study |
| **Spo2op1-DsRed** | Encodes spo2op1-DsRed (PA probe) | Nicolas Vitale |
| **Spo2op2-RFP** | Encodes spo2op2-RFP (PA probe) | Nicolas Vitale |
| **pRK5-HA** | CMV expression vector for N-terminal fusion of HA tag | (Martinez *et al.* 2016) |
| **pRK5-HA-ALFA** | CMV expression vector for N-terminal fusion of HA-ALFA tag | This study |
| **pRK5-HA-Vice** | CMV expression vector expressing *cbu2007* with N-terminal fusion of HA tag | This study |
| **pRK5-HA-ALFA-Vice** | CMV expression vector expressing *cbu2007* with N-terminal fusion of HA-ALFA tag | This study |
| **pRK5-HA-ALFA-Vice_1-213_** | CMV expression vector expressing *cbu2007***_1-213_** with N-terminal fusion of HA-ALFA tag | This study |
| **pRK5-HA-ALFA-Vice_214-395_** | CMV expression vector expressing *cbu2007***_214-395_** with N-terminal fusion of HA-ALFA tag | This study |
| **pRK5-HA-ALFA-Vice_214-345_** | CMV expression vector expressing *cbu2007***_214-345_** with N-terminal fusion of HA-ALFA tag | This study |
| **pLVX-GFP-N2** | CMV expression vector for C-terminal fusion of GFP | (Burette *et al.*, 2020) |
| **peGFP-RAB7** | Encodes human *rab7* with N-terminal fusion of GFP tag | Addgene #61803 Gia Voeltz |
| **pTO_CHMP3_LAP_EGFP** | Encodes human *chmp3* with N-terminal fusion of GFP tag | Addgene #101849 Daniel Gerlich |
| **pALIX_FLAG_GFP** | Encodes human *alix* with N-terminal fusion of GFP tag | Aurélien Roux (Larios *et al.* 2020) |
| **pTWIST_2007** | Encodes *6*x*HIS-cbu2007* codon optimized for expression in *E. coli* | Renaud Vincentelli |
| **pET28b_GST** | Encodes 6x*HIS-gst* | This study |
| **pGEX4T1** | Encodes GST | GE Healthcare |
| **pGEX4T1-RAB26** | Encodes GST-RAB26 | This study |
| **pGEX4T1-CHMP3** | Encodes GST-CHMP3 | This study |
| **pGEX4T1-ALIX** | Encodes GST-ALIX | Maryse Lebrun |
| **pCMV-NbALFA-mScarlet** | Encodes mScarlet-tagged nanobodies targeting ALFA | This study |
| **pCMV-NbALFA-mNeonGreen** | Encodes mNeonGreen-tagged nanobodies targeting ALFA | This study |
| **pmCherry-CHMP4B** | Encodes human *chmp4b* with C-terminal fusion of mCherry tag | Aurélien Roux (Larios *et al.* 2020) |
| **pCMVGag** | Encodes for HIV-1 Gag gene | Delphine Muriaux (Burniston *et al.* 1999) |
| **pLifeact-EGFP** | Encodes the GFP-tagged Lifeact peptide. | Delphine Muriaux |
| **lentiCas9-Blast** | Encodes Cas9 | Addgene #47948 |
| **pLentiGuide-Puro-CTRLsg1** | HIV1-based lentivector for stable expression of non-targeting sgRNA in mammalian cells | Siadous *et al.*, 2020 |
| **pLentiGuide-Puro-CTRLsg2** | HIV1-based lentivector for stable expression of non-targeting sgRNA in mammalian cells | Siadous *et al.*, 2020 |
| **Sanger lentiviral CRISPR vector ALIX-E11** | Lentivector for stable expression of ALIX-targeting sgRNA (GCTGCAATTTGGCTAGCTAAGG) in mammalian cells | This study |
| **Sanger lentiviral CRISPR vector ALIX-E12** | Lentivector for stable expression of ALIX-targeting sgRNA (GATGCCATCATAGCTAAATTGG) in mammalian cells | This study |
